# CATE: A fast and scalable CUDA implementation to conduct highly parallelized evolutionary tests on large scale genomic data

**DOI:** 10.1101/2023.01.31.526501

**Authors:** Deshan Perera, Elsa Reisenhofer, Said Hussein, Eve Higgins, Christian D. Huber, Quan Long

## Abstract

Statistical tests for molecular evolution provide quantifiable insights into the selection pressures that govern a genome’s evolution. Increasing sample sizes used for analysis leads to higher statistical power. However, this requires more computational nodes or longer computational time. CATE (CUDA Accelerated Testing of Evolution) is a computational solution to this problem comprised of two main innovations. The first is a file organization system coupled with a novel search algorithm and the second is a large-scale parallelization of algorithms using both GPU and CPU. CATE is capable of conducting evolutionary tests such as Tajima’s D, Fu and Li’s, and Fay and Wu’s test statistics, McDonald–Kreitman Neutrality Index, Fixation Index, and Extended Haplotype Homozygosity. CATE is magnitudes faster than standard tools with benchmarks estimating it being on average over 180 times faster. For instance, CATE processes all 54,849 human genes for all 22 autosomal chromosomes across the five super populations present in the 1000 Genomes Project in less than thirty minutes while counterpart software took 3.62 days. This proven framework has the potential to be adapted for GPU-accelerated large-scale parallel analyses of many evolutionary and genomic analyses.

## INTRODUCTION

To elucidate a population’s current genetic diversity, population genetics attempts to decipher the historic effects of selection pressures on genetic variation via an assortment of statistical tests (1–5). Such tests draw on genome-wide patterns of allele frequency and haplotype variation to discriminate neutral evolution from past instances of natural selection (6, 7).

Modern advancements in high throughput sequencing and digital simulations have resulted in an accruing volume of data. Upon completion of the human genome project in 2003 and the launch of Next Generation Sequencing (NGS) in 2005, the cost of genome sequencing dramatically dropped (8, 9). This major deviation from Moore’s law predictions has resulted in an unprecedented surge in genetic data resulting in repositories that span thousands of samples across tens of millions of markers to become commonplace (10). Such endeavours include the 1000 Genomes Project (11), 1001 Genomes Project (12, 13), gnomAD (14), and the UK biobank (15). The ability to process these large datasets in their entirety increases the power of the statistical tests resulting in more concrete conclusions (16). However, the ability to process these repositories in their entirety is strained by the requirement of massive amounts of computational resources in the form of memory and computing power (10, 17, 18).

To tackle this predicament, researchers need solutions conducting genome wide calculations on expansive repositories in a fast, resource-efficient, and scalable nature (10, 17, 19). However, popular computational tools, though capable of being executed on HPC (High Performance Computing) clusters, have been designed with the goal of processing data that spans a few hundred individuals across thousands of polymorphic sites (20–22). They use single thread algorithms that do not exploit the prevalent parallel processing technologies made available through multi core CPUs (Control Processing Unit), and GPUs (Graphical Processing Unit) (10, 21, 23, 24).

The parallelization of algorithms has been proven to reduce computational time (25–27), and most evolutionary tests can be parallelized due to their nature of independently evaluating polymorphisms that are in different genes or regions of the genome. If catered to, algorithms can be designed to harness not only the CPU’s parallel processing capabilities with tens of cores but also the large-scale parallelization capabilities of GPUs which have thousands of cores. With 2012’s stable release of NVIDIA’s CUDA (Compute Unified Device Architecture) framework, which provides an interface for programming the GPU, programmable GPUs are steadily becoming commonplace in both general-purpose and specialized computing (26). If organized properly, data retrieval from storage can similarly be parallelized (16, 28).

Here, we present a solution, CATE (CUDA Accelerated Testing of Evolution), algorithmically optimized to synergistically harness the computer’s parallel processing technologies of the GPU, CPU, and SSD (Solid State Drive) to process expansive variant call data. We have built a scalable program using NVIDIA’s CUDA platform together with an exclusive file hierarchy, coupled with our own novel search algorithm called CIS (Compound Interpolated Search) to conduct six different tests frequently used in molecular evolution: Tajima’s D, Fu and Li’s D, D*, F and F*, Fay and Wu’s H and E, McDonald–Kreitman test, Fixation Index, and Extended Haplotype Homozygosity (EHH).

The techniques implemented in CATE enable us to overcome hurdles that plague the processing of large data repositories such as sequential file access and requirements for substantial amounts of expensive onboard memory. For instance, CATE’s unique file hierarchy allows random access to the file system and the bespoke Compound Interpolated Search (CIS) search algorithm enhances this endeavour.

Our solutions are not limited to the computation of statistical tests for selection. The fragmentation of large files along with the use of our CIS search algorithm improve the efficiency of data retrieval and reduce stresses on onboard memory, and the use of GPU systems for large scale parallelization cut down on processing times. Our algorithmic solutions can act as a template to be implemented in other software systems designed for large-scale parallel processing of genomic data via the GPU.

CATE is designed to conduct large-scale parallel processing of segregating sites using the GPU. Our solution is not only magnitudes faster than existing tools, but it is both scalable and resource efficient. The CATE software’s flexibility enables it to be implemented on both personal computers equipped with CUDA enabled GPUs and HPCs.

## MATERIALS AND METHODS

### Innovations

Evolutionary tests require the collection of allelic data from genomic regions across populations. In large datasets the process of finding and collecting the data from the regions of interest can be time consuming. This is because these datasets are organised into a singular file forcing sequential access spanning thousands to even millions of lines. The expansiveness of these datasets leads to latencies in processing time. CATE attempts to solve the problem of latency in conducting evolutionary tests through two key innovations: a unique file hierarchy together with a novel search algorithm (CIS) and GPU level parallelisation with the Prometheus mode (**Supplementary materials Section 2**). Each of these innovations address specific drawbacks faced with big data genomic analysis. Together, they facilitate CATE to have a high rate of computation and resource efficiency.

CATE also has a built-in resume feature (**Supplementary materials Section 2.5**). If the program abruptly ends before the completion of an evolutionary test function, it can return to its last successfully completed query. It will then continue the process from its last stop point. We have designed this feature to be seamless so that the user simply needs to execute CATE without any modifications. CATE is capable of recognising that the current query set was previously executed, which triggers the resume feature.

### Software overview

CATE, abbreviated from CUDA Accelerated Testing of Evolution, is a command line interface software written in C, C++, and CUDA. It is a Linux and Unix compatible free and open-source tool available at the GitHub repository https://github.com/theLongLab/CATE. CATE is under the MIT license. The repository is equipped with the source code, complete with its documentation, examples of execution, and a user guide (**Supplementary materials Section 1.1**).

CATE’s general architecture (**Figure 1**) is designed to run on almost any system equipped with a CUDA-enabled GPU with minimal input from the end user (**Supplementary materials Section 1.1**). The program will first verify the input function, configure the intermediate and output folders, index the file hierarchy system, and will then initiate the processing of the query regions. The general architecture guarantees the efficient utilisation of hardware resources, such as memory and processing power resulting in a robust and lightweight tool.

**Figure 1.**
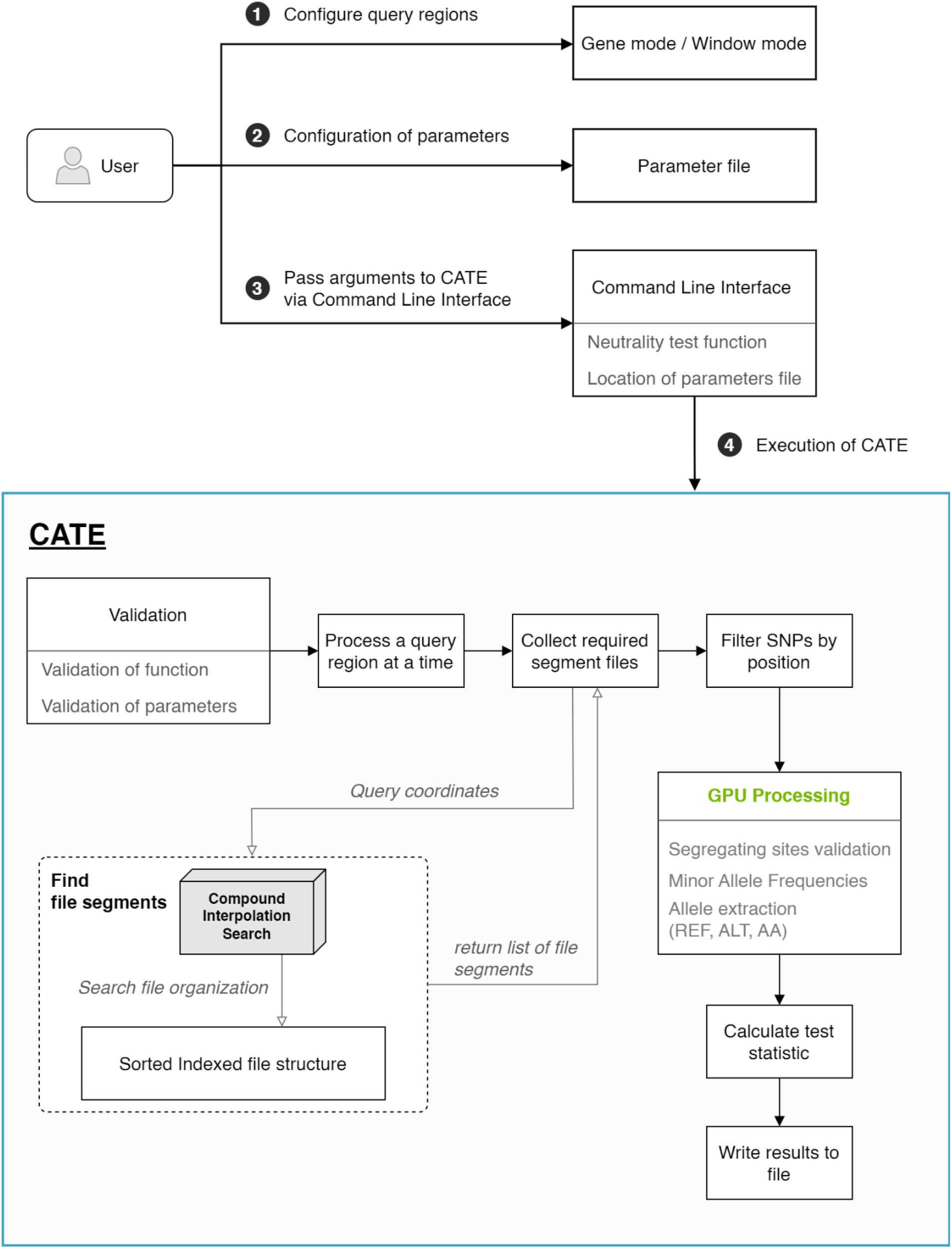
Overview of the general workflow of CATE. The user will begin by first defining the mode of analysis either by gene regions or by fixed windows of analysis. If the user prefers a window wise analysis the window size and step size will be included in the parameters file. The user will secondly configure the parameters file which will enable CATE to locate the output folders to print its results as well as the location of the segmented file structure. Thirdly the user will pass arguments via the Command Line Interface (CLI) and fourth execute CATE. CATE will conduct analysis by identifying the file segments that will contain the required SNP data. The SNPs that satisfy the query region being processed will be within these file segments. It will then collect and filter this data by position to obtain only the necessary SNPs. These SNPs will be processed together by parallelisation in the GPU. Finally using the extracted information CATE will calculate the test statistic for the user specified neutrality tests and write the results to an output file. The figure depicts CATE on its normal mode where it processes one query region at a time. This process will be repeated till all query regions are processed.

CATE provides six tests of neutrality with an additional eight complementary tools for genomic data processing including the tool to automatically create our unique segmented file structure (**Supplementary materials Section 1.2**). These tools are designed to help the users conduct utilitarian tasks such as data extraction and merging on different genomic data files such as FASTA and General Feature Format (GFF). CATE is also equipped with a function to extract haplotype information from a query region.

All main functions of CATE are dependent on the availability of a CUDA enabled GPU. These GPUs are proprietary to the NVIDIA cooperation (27). To interact with CATE (**Figure 1** and **Supplementary materials Section 1.1**) the end user will pass functions via the command line and configure the parameters of these functions using a tailored parameter file written in JSON format (**Supplementary materials Figure S – 1 and Section 1.3**).

For four out of the six tests, CATE provides both a gene file mode and a sliding window mode. These tests include Tajima’s D, Fu and Li’s D, D*, F and F*, Fay and Wu’s H and E, and Fixation Index. The remaining two tests of McDonald Kreitman and EHH deviates from the aforementioned four tests in their mechanism of analysis. McDonald Kreitman analyses a gene’s coding region known as an Open Reading Frame (ORF) and EHH requires the specification of a core haplotype. There is also an option to conduct the first three neutrality tests of Tajima’s D, Fu and Li’s D, D*, F and F*, Fay and Wu’s H and E together at once, called Neutrality full. Gene file mode allows the user to configure a series of independent regions to be analyzed. Window mode is the window-wise analysis technique available in most evolutionary testing software. CATE can be configured to analyse overlapping and non-overlapping windows as well as continuous sliding windows. (10, 21, 29).

For its neutrality tests, CATE comes equipped with two performance modes, a normal mode and a high-performance mode called *Prometheus*. Users will be able to configure core usage, SSD access, and memory control via this feature to allow CATE to achieve maximum use of the hardware (**Figure 3**).

**Figure 2.**
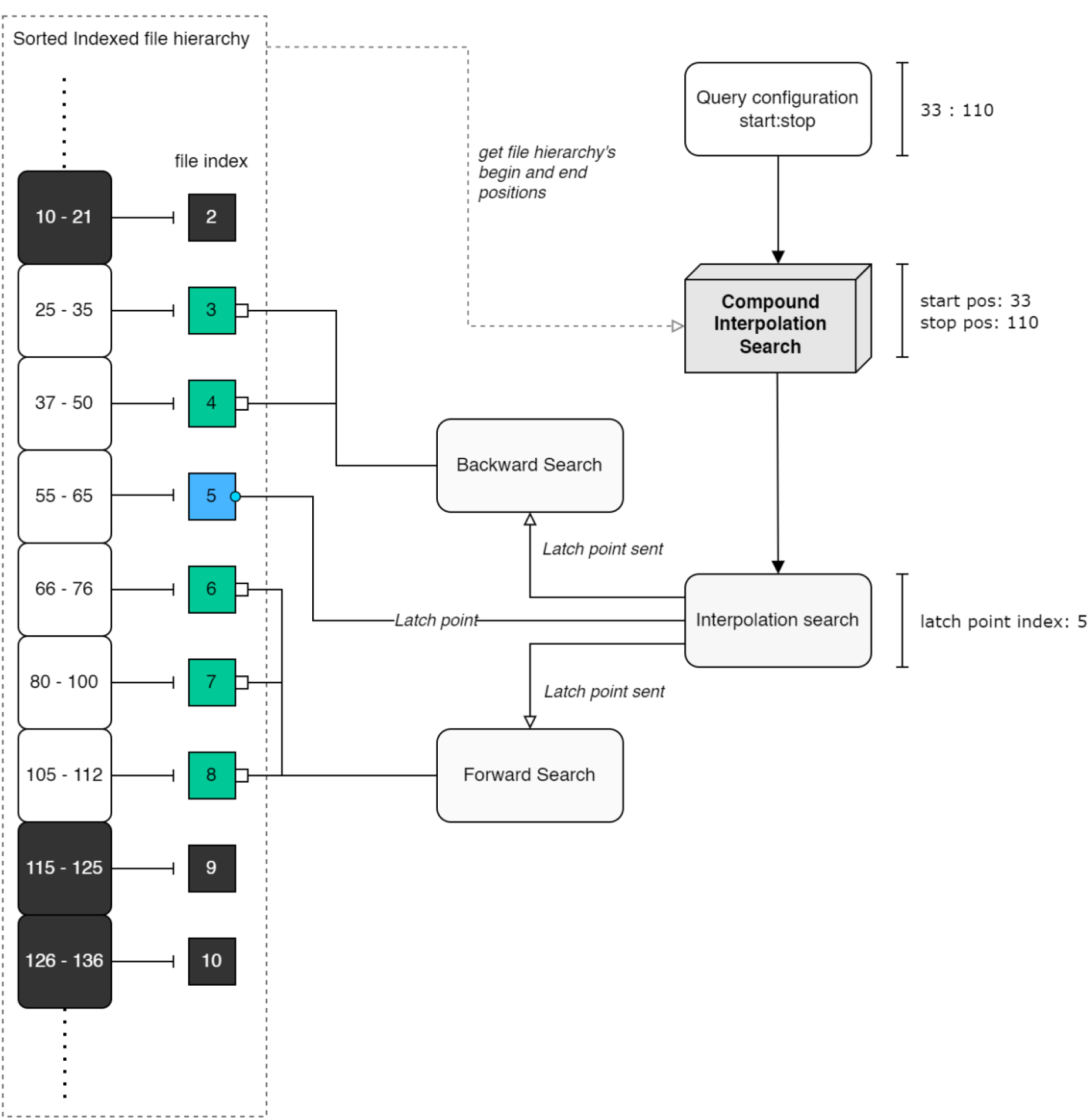
Illustration of the Compound Interpolated Search (CIS) algorithm’s architecture with an example. It collects required segment files that satisfy the query region of interest. The data that satisfies the query region lies within these segment files. The algorithm uses three CPU threads to complete its function. The interpolation search algorithm is used to identify the latch point. The latch point is the first file segment that satisfies the query region. The algorithm will search for the remaining segment files by searching the space around the latch point using two independent sequential searches. Due to their independent nature, they can be executed in parallel.

**Figure 3.**
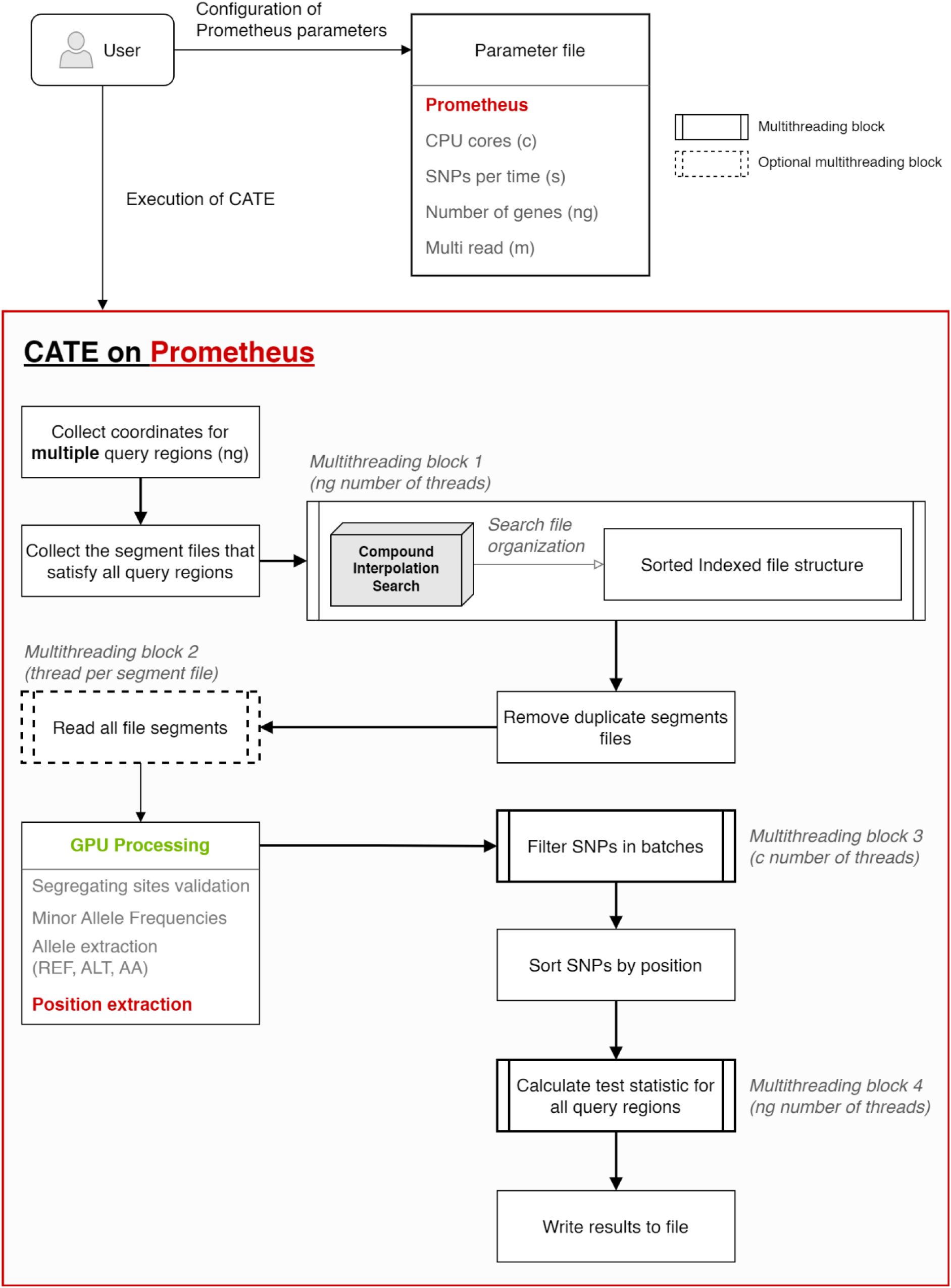
Diagrammatic overview of CATE’s high performance mode Prometheus. Compared to CATE’s normal mode the user configures Prometheus using four additional parameters in the parameters file. Once CATE is executed it will detect the activation of Prometheus. In contrast to its normal mode, Prometheus processes multiple query regions together as a batch. It uses four additional parallel processing instances to achieve this. They are depicted as Multithreading blocks, with the number of threads used by each presented in brackets. Multithreading block 1 is used to determine the specific file segments required to satisfy each query region, followed by the removal of redundant file segments. Multithreading block 2 is optional. It is activated in the presence of SSD drives. If present all segment files will be read at once, if absent each file will be read sequentially. To prevent the overloading of GPU memory the number of SNPs the GPU handles at a time is controlled by the parameter SNPs per time (s). The Multithreading block 3 will then process SNPs and validate them as segregating sites in batches. Once the SNPs are sorted by position the respective neutrality test statistics will be calculated in parallel for all query regions in the Multithreading block 4.

### File structure and Compound Interpolated Search (CIS) algorithm

Our file organization approach addresses and overcomes a major drawback of sequential access in current VCFs. CATE creates its file hierarchy by splitting a single VCF file into smaller segments. Each segment has a fixed number of variants. These segments are then labelled by the range of the variant sites’ base pair positions that are stored within them (**Supplementary materials Section 2.1**). This allows us to sort the file segments by base pair position enabling it to be searched using quick search algorithms such as binary search and interpolated search (30, 31).

CATE’s built-in tool VCF Splitter is designed for this exercise of segmentation. If needed VCF Splitter can be configured to filter the VCFs by population, MAF (Minor Allele Frequency), and allele counts as it creates the file segments. This organised and sorted file structure enables random accessing of the dataset in a quick and efficient manner. The CIS algorithm (**Figure 2**) identifies the segments that satisfy the user’s query. CATE will then directly read these segments and extract the relevant data. This saves time as the alternative would require the software to iterate over the entire VCF file before reaching the required target region of interest.

The CIS algorithm consists of two search algorithms: interpolated search and sequential search (**Supplementary materials Sections 2.2**). They are configured to work together to find the collection of file segments that satisfy the query region. The CIS algorithm operates by taking in the query region’s range. Then using the interpolation search, it finds a segment from the file hierarchy that satisfies the query region. We call this the *latch point*. The CIS algorithm then produces two sequential search algorithms in separate threads around this latch point to search for the rest of the segment files. Starting from the latch point one sequential search algorithm goes forward in the sorted segment files hierarchy and the other goes backwards in the file hierarchy. In this manner, all the segments that satisfy the query region are collected. The relevant SNPs that satisfy the query region is nested within the list of segments files found by the CIS algorithm. These files will then be read in a sequential manner validating each position to be within the (nested) target query region’s range.

Our CIS algorithm enables us to parallelise the search function. The adaptation of the interpolated search algorithm enables it to intelligently find the starting point of the search space. This search strategy enables keeping the time for data retrieval and the subsequent processing speeds constant regardless of the positioning of the query region in the chromosome. Additionally, since the file segments are separate, the file structure enables the accessing of multiple files at the same time through SSD technologies. As far as we know CATE is the first software implementation in the genomic space with a framework capable of using SSD technologies to conduct simultaneous access of multiple files. SSDs can handle multiple read-and-write instructions in contrast to their single HDD (Hard Disk Drive) counterparts which can only perform a single read or write function at a time (28, 32).

### GPU level parallelization for evolutionary tests and the Prometheus mode

Parallelisation is done by assigning each GPU thread a single SNP. The thread knows which region of the array to process in relation to the SNP it was assigned by referring to the respective start and stop positions. This is the main algorithmic implementation that enables large-scale parallel processing of genomic data. A single query region can contain anywhere from a few hundred to hundreds of thousands of SNPs and these SNPs span over multiple individuals. The GPU architecture with thousands of thread counts is ideally suited for this work. The GPU can be parallelised to extract relevant information such as the Minor Allele, its Frequency (MAF), the Ancestral Allele (AA) information and more from all SNPs in unison.

Additionally, a series of bespoke optimizations per evolutionary test were performed (**Supplementary materials Section 3**). This allowed the optimisation of the speed and resource efficiency of each function. These developments include preventing redundancy in data reads in the F_st_ algorithm (**Supplementary materials Section 3.2**) when handling multiple populations, augmentation of the EHH algorithm to have the least amount of redundancy when searching for unique haplotypes, and finally the use of the GPU in the MK test function when finding ORFs and in the determination of substitution types.

### Prometheus: High-performance mode’s architecture

The neutrality tests of Tajima’s D, Fu and Li’s D, D*, F and F*, Fay and Wu’s H and E treat each segregating site independently. This characteristic can be exploited to enable further parallelization, leading to faster computation and better resource usage. To address this, a separate high-performance mode for called Prometheus was designed (**Supplementary materials Section 2.4**).

CATE’s Prometheus has a slightly different architecture (**Figure 3**) from its normal mode. The Prometheus architecture focuses mainly on batch processing of multiple query regions at the same time, whereas in normal mode CATE will process only a single query region at a time. To accommodate for this capability Prometheus is equipped with a series of multithreading blocks. These multithreading blocks as depicted in **Figure 3** focus on handling batches of data in parallel thereby reducing time otherwise spent on sequential processing of the data by a single thread or process. These multithreading blocks make maximum use of the computer’s parallel processing capabilities, expanding from beyond the GPU to the CPU (multithreading blocks 1, 3, and 4 in **Figure 3**) and even the SSD (multithreading block 2 in **Figure 3**).

To activate Prometheus the user must define four additional parameters: the number of CPU cores that can be used at a time, the number of SNPs the GPU can process at a time, if multi-thread is available for SSD file access, and finally the number of query regions to be processed together (**Supplementary materials Table S – 2** and **Figure S – 4**).

CATE will begin by collecting the start and stop positions of multiple query regions. It will then determine the unique list of file segments required to satisfy this collection of query regions using the CIS algorithm and proceed to read them. If the hardware is equipped with an SSD or flash storage multiple file segments will be read at the same time. The collected SNPs will then be processed in the GPU. The extracted SNPs’ information will then be used during the processing of each query region which is conducted in parallel by CPU multithreading (**Supplementary materials Section 2.4**).

Prometheus becomes faster than CATE’s normal mode because it saves a lot of time that is otherwise lost for redundant file reads and SNP processing. Prometheus achieves this leverage because it needs to process SNPs that can satisfy multiple query regions only once due to its batch processing strategy. Additionally, by distributing the query regions across multiple CPUs Prometheus can better parallelise the batch processing task as well, further improving its efficiency.

This exclusive architecture of Prometheus enables the user to not only customize the resource availability of CATE but allows the software to make maximum use of the resources at its disposal. The optimisation of the Prometheus parameters will be dependent on the user’s computer hardware and the size of the data repository. For instance, a user with more memory in their GPU and RAM can enable Prometheus to process a higher number of query regions at a time, and inversely if the repository has a large sample size, they can reduce the number of SNPs being processed concurrently by the GPU. This strategy of batch processing allows the orchestration of the program code, data, and architecture to work together. It also enables CATE to use all multiprocessing hardware available in modern computing which includes CPUs, GPUs, and SSDs.

### Benchmarking

CATE is equipped with six evolutionary tests spread across eight functions. To determine the robustness of CATE we have tested it against three expansive data sets namely, the 1000 Genomes dataset, the 1001 Genomes dataset, and SARS-CoV-2 data from GISAID (**Table 1**). These repositories were selected for their large sample sizes and curated polymorphic data (**Supplementary materials Section 4**).

**Table 1.** Details of candidate data repositories used to assess CATE’s performance. The datasets were selected for their vast collection of polymorphisms and samples. CATE was tested against these repositories to assess its limitations.

| Repository | Ploidy | Sample size | Chromosomes | SNPs |
| --- | --- | --- | --- | --- |
| 1000 Genomes Project | 2 | 3115 | 22 | 77,818,345 |
| 1001 Genomes Project | 2 | 1135 | 5 | 375,146 |
| GISAID (sample dataset) | 1 | 5081 | 1 (NC_045512.2) | 4,789 |

CATE’s segmented file structure is a crucial component of its functionality that enables its speed and resource efficiency. Therefore, it is important that it can be created in a fast and efficient manner. The time taken to create the segmented file structure for over 81 million SNPs spread across the 22 autosomal chromosomes of the 1000 Genomes data was assessed for this purpose. To ensure robust testing we made CATE create file structures with and without filtration of SNPs by MAF. MAF filtration was set for common SNP at above 0.05 (33).

CATE was benchmarked for each test alongside existing software tools (**Table 2**). This enabled us to assess its speed and accuracy. The evolutionary test results of the tools were contrasted against that of CATE’s outputs, and the total run times and resource usage were also evaluated. CATE’s primary design objective is to be quicker, scalable, and more resource efficient. In all instances, it was ensured to make the benchmarking comparable by maintaining equivalent hardware and datasets.

**Table 2.** Description of the assessed evolutionary tests available on CATE and the tools against which it was tested. CATE was tested against complementary software where possible and in instances where complimentary software was not available a controlled example was used.

| Evolutionary test | Complementary software | Repository |
| --- | --- | --- |
| Tajima's D | PopGenome, vcf-kit | 1000 Genomes Project, 1001 Genomes Project, GISAID |
| Fu, and Li's D, D*, F and F* | PopGenome | 1000 Genomes Project, 1001 Genomes Project, GISAID |
| Fay and Wu's normalized H and E | PopGenome | 1000 Genomes Project, 1001 Genomes Project, GISAID |
| F<sub>st</sub> | NA | 1000 Genomes Project, 1001 Genomes Project |
| EHH | selscan | 1000 Genomes Project, 1001 Genomes Project |
| McDonald–Kreitman | NA | 1000 Genomes Project |

CATE was used to process each dataset under three different test types (**Table 3**). The three neutrality tests by Tajima, Fu, and Li and, Fay and Wu were subjected to three different test types (**Supplementary materials Section 4.1, Section 4.2,** and **Section 4.3**). The “all genes” test type tested all known gene regions in a genome. For this test, all the available genes’ coordinates were downloaded using the BioMart R package (34). The coordinates for Genome Reference Consortium Human genome build 37 (GRCh37) was used. This build was used because Phase 3 of the 1000 Genomes Project used the GRCh37 reference genome to map the variants (35).

**Table 3.** Details of the tests that were conducted using each data repository. The array of varying test types was conducted to understand the flexibility and limitations of CATE, in terms of resource usage and processing time. The neutrality tests of Tajima’s D, Fu, and Li and, Fay and Wu statistics were also tested against CATES Prometheus mode as well. Window mode requires the window size and step size parameters to be configured in base pairs (bp) and Sliding window mode the step size is a single Nucleotide Polymorphism (SNP).

| Repository | Test type | Evolutionary test | Parameters |  | Query regions |
| --- | --- | --- | --- | --- | --- |
| 1000 Genomes Project | All genes | Tajima's D<br>Fu, and Li's D, D* F and F*<br>Fay and Wu's normalized H and E<br>F<sub>st</sub><br>EHH<br>McDonald–Kreitman | NA |  | 54,849 |
|  | Window | Tajima's D<br>Fu, and Li's D, D*, F and F*<br>Fay and Wu's normalized H and E<br>F<sub>st</sub> | Window size | 10,000 bp | 279,621 |
|  |  |  | Step size | 10,000 bp |  |
|  | Sliding window | Tajima's D<br>Fu, and Li's D, D*, F and F*<br>Fay and Wu's normalized H and E | Window size | 10,000 bp | 77,572,883 |
|  |  |  | Step size | 1 SNP |  |
|  | SNP<br>BP | EHH | SNP window<br>BP window | 250 SNPs<br>100,000 bp | 11,000 |
| 1001 Genomes Project | All genes | Tajima's D<br>Fu, and Li's D, D* F and F*<br>Fay and Wu's normalized H and E<br>F<sub>st</sub><br>EHH<br>McDonald–Kreitman | NA |  | 28,495 |
|  | Window | Tajima's D<br>Fu, and Li's D, D*, F and F*<br>Fay and Wu's normalized H and E<br>F<sub>st</sub> | Window size | 10,000 bp | 11,912 |
|  |  |  | Step size | 10,000 bp |  |
|  | Sliding window | Tajima's D<br>Fu, and Li's D, D*, F and F*<br>Fay and Wu's normalized H and E | Window size | 10,000 bp | 384,786 |
|  |  |  | Step size | 1 SNP |  |
|  | SNP<br>BP | EHH | SNP window<br>BP window | 250 SNPs<br>100,000 bp | 2,500 |
| GISAID SARS-CoV-2 | All genes | Tajima's D<br>Fu, and Li's D, D* F and F*<br>Fay and Wu's normalized H and E | NA |  | 11 |

The “window” test type conducted neutrality tests of specific regions along the chromosome and finally the “sliding window” moved across each SNP in the chromosome and calculate a fixed window of SNPs from that position. All tests were subjected to the “all genes” test type and EHH has two unique options of “SNP” and “BP” which allows the user to calculate the degradation of the EHH value around a SNP (**Supplementary materials Section 3.3**) (36, 37).

The hardware used to test CATE is two-fold, both configurations were equipped with CUDA enabled GPU’s. The first configuration was used to provide equal capabilities to both CATE and the software it was tested against. The second configuration was a high-end platform named Narval by Compute Canada. Narval was used because Prometheus required a platform with SSD capabilities ensuring that all its features could be tested. Additionally, this platform was used to benchmark the maximum speed capabilities of CATE given its state-of-the-art resources. The first configuration consisted of Tesla V100-PCIe-16GB GPUs, 2x Intel(R) Xeon(R) Gold 6148 CPU with 2.40GHz (Skylake, 2019), and HDD storage capabilities. The second configuration was a higher-end system by Compute Canada comprising of NVidia A100-40GB GPUs, 2x AMD Milan 7413 CPU with 2.65 GHz 128M cache L3 and SSD storage. By using two separate hardware platforms we were also able to evaluate the changes in the efficiency of CATE in relation to different hardware.

## RESULTS

To illustrate the efficiency of CATE a series of benchmark tests to assess its speed and accuracy were conducted. Each evolutionary test function was assessed for its performance against various datasets and mainstream computational tools. CATE comes complete with three separate neutrality tests, each with its own set of statistics creating a total of seven separate values. Additionally, the Neutrality test functions are equipped with CATE’s high-performance mode Prometheus. We tested both these modes extensively.

### Generation of segmented file structure

The efficiency of creating CATE’s file structure was assessed via the 22 autosomal chromosomes from the 1000 Genomes Project. The results of the test are depicted in **Table 4** and a detailed breakdown of the per chromosome run times are depicted in **Supplementary materials Table S – 11**.

**Table 4.** Summary overview of the time taken to create CATE’s segmented file structure for all 22 autosomal chromosomes from the 1000 Genomes dataset. We tested the time taken to create file structures both in the presence and absence of a MAF filter.

| Total SNPs | Average SNPs | MAF filter | Total run time (s) | Average run time (s) |
| --- | --- | --- | --- | --- |
| 81,271,767 | 3,694,171.23 | NA | 14644 | 665.64 |
|  |  | Greater than 0.05 | 8365 | 380.23 |

CATE had a significantly faster run time of 6.34 minutes on average per chromosome in the presence of a MAF filter in contrast to the absence of a filter which averaged 11.09 minutes per chromosome. We believe that this may be caused by the increased number of SNPs that have to be written to the hard disk in the absence of a MAF filter.

### Frequency Spectrum Neutrality tests

Neutrality tests were validated by estimating the values of CATE with two complementary software vcf-kit and PopGenome. PopGenome enabled the validation of all three neutrality tests and vcf-kit was limited to only Tajima’s D (10, 21). CATE’s results were generally identical to both software (**Supplementary materials Tables S – 12, S – 13,** and **S – 14**).

We then estimated the speed and resource usage of CATE to conduct the neutrality tests on large scale data sets. For this, the analysis was conducted on all 22 autosomal chromosomes available in the 1000 Genomes Project. Initial tests showed that vcf-kit was unable to process the entire chromosome-wide VCF files and faced memory issues. Furthermore, it was only capable of conducting window-wise analyses. PopGenome proved to be able to process the data in both gene-wise and window-wise test types. PopGenome conducts all three neutrality tests at once. This test type was comparable to CATE’s Neutrality full function.

CATE in its normal mode alone was significantly faster than PopGenome. To process a total of 54,849 query regions across 22 chromosomes CATE was 22.89 times faster than PopGenome (**Figure 4** and **Supplementary Table S – 15**). When Prometheus was activated, this performance was further accelerated. CATE on Prometheus was 7.95 times faster than CATE on normal mode and 182.01 times faster than PopGenome. CATE averaged a time of ten minutes to process the entire collection of genetic regions of each chromosome’s variant call data on its normal mode and about only 1.30 minutes when Prometheus was activated, whereas PopGenome averaged 3.95 hours per chromosome. (**Supplementary Table S – 15**).

**Figure 4.**
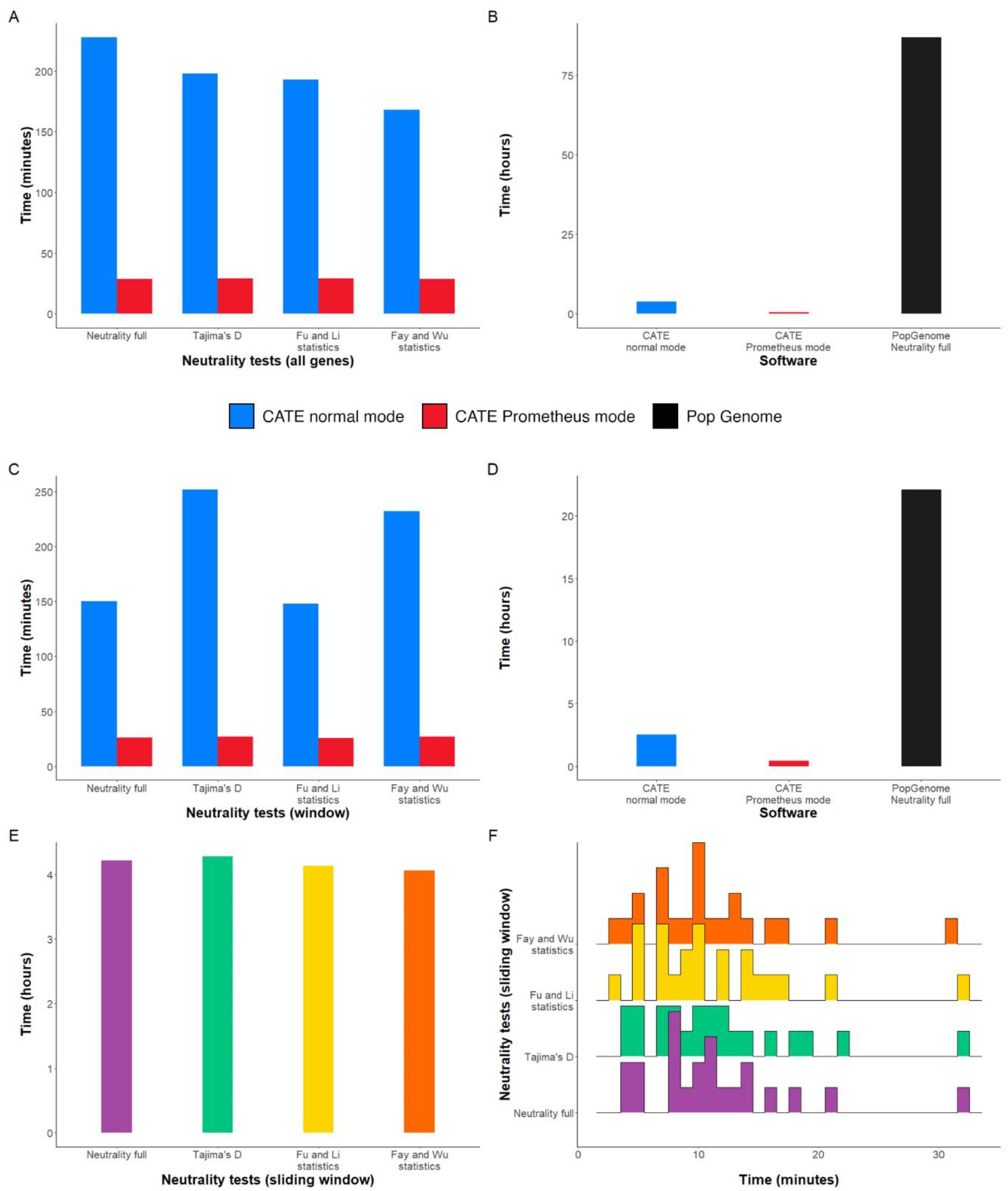
Assessment of CATE’s performance by comparing the time taken to conduct the four neutrality tests (Tajima’s D, Fu and Li’s, and Fay and Wu’s test statistics) on 22 autosomal chromosomes from the 1000 Genomes dataset. (**A**) CATE’s normal mode vs. CATE’s Prometheus high performance mode for the all-genes test type. (**B**) CATE’s normal mode vs. CATE’s Prometheus vs. Pop Genome for analysing all three neutrality tests using the Neutrality full function for the all-genes test type. (**C**) CATE’s normal mode vs. CATE’s Prometheus high performance mode for the window test type (window size = 10,000, step size = 10,000). (**D**) CATE’s normal mode vs. CATE’s Prometheus vs. Pop Genome for analysing neutrality tests using the Neutrality full function for the window test type (window size = 10,000, step size = 10,000). In all instances CATE outperforms PopGenome in processing time with increased speed when Prometheus is activated. (**E**) Bar plot of Prometheus’ total run times for the sliding window analysis (window size = 10,000, step size = 1 SNP) resulting in a total of 77 million SNPs per neutrality test. (**F**) Distribution plot breaking down Prometheus’ run times to process each autosomal chromosome during the sliding window analysis. It shows that on average most chromosomes were processed within 9 to 12 minutes. (**A – E**) Bar plots show the total time taken by each software to process all 22 chromosomes per neutrality test.

When processing window wise, which comprised 279,621 query regions, CATE on normal mode was 8.83 times faster than PopGenome (**Figure 4** and **Supplementary Table S – 15**). Software such as PopGenome are designed for the window-based calculation method, which explains why it performed better on the window wise analysis than the all genes test type. When Prometheus was activated, CATE was 51.44 times faster than PopGenome. CATE averaged a time of 6.82 minutes to process the variant call data spanning across each chromosome on its normal mode and about only 1.17 minutes when Prometheus was activated whereas PopGenome averaged about one hour per chromosome (**Supplementary Table S – 15**).

To further test the performance of CATE’s Prometheus we conducted a Sliding window analysis of the entire 1000 Genome project’s 22 autosomal chromosomes (**Figure 4** and **Supplementary Table S – 15)**. In the Sliding window analysis, the step size is the incrementation from one SNP to the next. We used a fixed window size of 10,000 bp. This was a total of 77,572,883 query regions. CATE conducted the complete analysis in 4.21 hours with an average time of 32.34 minutes per chromosome (**Supplementary Table S – 15**). This meant that CATE was able to process over 77 million SNPs across 22 chromosomes in less time than PopGenome took to process a single chromosome containing about 5000 query regions.

Both the window and sliding window test types enabled the assessment of sequential accessing of the variant call file structure of CATE against software such as PopGenome and vcf-kit. The gene-wise analysis allowed the testing of the random accessing capabilities of CATE. CATE was specifically designed for this use case scenario and its catered optimizations enabled it to utilize its capabilities to gain leaps in performance.

After the successful assessment of CATE’s speed, accuracy, and efficiency through the 1000 Genomes Project dataset, we tested for its robustness using the 1001 Genomes Project and the GISAID SARS-CoV-2 datasets. Despite the 1001 Genome Project variant call data having regions of fragmented and incomplete reads (38) CATE was able to process the data effectively. In all genes mode CATE completed the Neutrality full function for a total of 25,495 genes spread across 5 chromosomes of *Arabidopsis thaliana* in under 200 seconds averaging 40.00 seconds per chromosome. Keeping similar parameters to the 1000 Genomes Project’s Window and Sliding Window analyses (**Table 3**) CATE neutrality full function was conducted on the 1001 Genomes Project data. The Window mode was completed in 96 seconds for a total of 11,912 query regions and the sliding window analysis was completed in 39.25 minutes for a total of 384,786 query regions.

### Fixation Index (F_st_)

The population wide F_st_ function was validated for accuracy using a model example (**Supplementary materials Section 5.3**). The function was benchmarked for both the window and all gene test types. F_st_ was calculated for a combination of three subpopulations of African (AFR), East Asians (EAS), and Europeans (EUR) from the 1000 Genomes Project (**Supplementary materials Table S – 20**) and for the 1001 Genomes project a combination of the three subpopulations of Asia, Western Europe, and Central Europe was used (**Supplementary materials Table S – 21**). The summary results are depicted in **Table 5**.

**Table 5.** Summary overview of the total run time taken for each test and the average run time per chromosome from the 1000 Genomes Project for the Fixation Index function. The results below depict not only CATE’s ability to process a large number of query regions but that it can navigate through multiple subpopulations of data to generate its results. In this instance, a combination of three subpopulations was used. The window test type consisted of both the window size and step size being 10,000 bases.

| Repository | Test type | Query regions | Total run time (s) | Average run time (s) |
| --- | --- | --- | --- | --- |
| 1000 Genomes Project | Window | 279,621 | 11,873 | 539.68 |
|  | All genes | 54,849 | 6,037 | 274.41 |
| 1001 Genomes Project | Window | 11,912 | 112 | 22.40 |
|  | All genes | 28,495 | 252 | 50.40 |

CATE averaged run times of 8.99 minutes and 4.57 minutes per chromosome from the 1000 Genomes Project for the window and all genes test types respectively. It was significantly quicker with the smaller dataset of 1001 Genomes with times of 50.40 seconds and 22.40 seconds for the all genes and window analyses.

### Extended Haplotype Homozygosity (EHH)

CATE has two versions of the EHH test spread across four modes, the SNP/ BP mode, and the FIXED/ FILE mode. Both were tested for (**Supplementary materials Section 3.3** and **Section 5.4**). The results of our EHH function were validated using the selscan software (17). The results of CATE were the same as that of selscan’s (**Supplementary materials Table S – 22** and **Table S – 23**). The benchmark times of the EHH functions are shown in **Table 6** (**Supplementary materials Table S – 24** and **Table S – 25**).

**Table 6.** Summary overview of the EHH functions two test types. The all genes test type requires the function to discover the unique haplotypes that may be present in the core region of interest, and then calculate the EHH value for each haplotype separately.

| Repository | Test type | Query regions | Average number of core haplotypes | Average run time (s) |
| --- | --- | --- | --- | --- |
| 1000 Genomes Project | SNP | 11,000 | Core haplotypes reference allele (0) and alternate allele (1) | 2,063.36 |
|  | BP |  |  | 3,815.73 |
|  | All genes | 54,849 | 13,574.68 | 107.82 |
| 1001 Genomes Project | SNP | 2,500 | Core haplotypes reference allele (0) and alternate allele (1) | 388.40 |
|  | BP |  |  | 330.80 |
|  | All genes | 28,495 | 44,430.80 | 50.40 |

In SNP mode CATE was assigned to explore a region of 250 SNPs both forward and backward in space from the core SNP creating a search window of 500 SNPs around the core SNP. The total analysis of took on average about 34.39 minutes per chromosome in the 1000 Genomes Project dataset and about 6.47 minutes on average per chromosome for the 1001 Genomes Project dataset. In both instances, a chromosome had 500 randomly selected core SNPs as the query regions. For the same dataset in the BP mode which explored a 200,000 bp window (100,000 bp on either side of the core SNP) around the core SNP the times were about 1 hour per chromosome for the 1000 Genomes Project data and about 5.51 minutes when processing the 1001 Genomes Project data.

In all genes test type analysis for a specified core and extended region (Extended region spanned from the start of the core region’s position to 10,000 base pairs forward in position) was done. For the 1000 Genomes Project CATE was able to complete the test with an average time of 1.8 minutes per chromosome with each chromosome having about 13,574 unique core haplotypes. Therefore, a chronological analysis of each chromosome at a time would take less than an hour (39.53 minutes). For the 1001 Genomes Project data CATE averaged 50.40 seconds per chromosome in the all genes analysis with each chromosome having about 44,430 unique core haplotypes on average.

### McDonald–Kreitman (MK) Neutrality Index (NI)

The MK test function was validated using a model example (**Supplementary materials Section 5.5**). Due to the complexities of setting up multiple genetic alignments and ORF validations required for the MK test, we conducted a benchmark test on chromosome one from the 1000 Genome dataset. An ORF is found in a region of DNA that spans between a start and stop codon consisting entirely of triplets with each triplet coding for an amino acid. A valid ORF would translate into an amino acid sequence that is responsible for a specific protein (39). We essentially performed a stress test on CATE, making the software execute the maximum number of calculations under the most unfavourable circumstances. We did an all genes test, for a chromosome-wide alignment of chromosome 1 of the outgroup species *Pan troglodytes* (chimpanzees) (NCBI Accession No. CM000314.3). CATE was set to find ORFs on its own for the entirety of the gene set. The analysis was completed in 2.01 hours. Additionally, CATE was conducting the analysis on a sample size of 2,504 individuals over the entire chromosome’s length.

### Hardware requirements

On all tests CATE (without Prometheus) required a CPU usage ranging from 29.2% to a maximum of 94.2% in a configuration of 5 CPU cores. CATE was able to run on an average of 30% CPU usage for all tests except for the McDonald Kreitman test for which it required about 90% of CPU usage on average. Between 3.73 Gb and 15.00 Gb of RAM for all six neutrality test functions was used. On the GPU CATE used a maximum of 381.00 Mb of memory and the GPU usage was between 2% to 10% capacity.

In comparison, to conduct the three neutrality test statistics for Tajima, Fu, and Li and, Fay and Wu using PopGenome it required between 93.0% to 97.1% of single core CPU usage with RAM usage lying between 12.00 Gb to 25.57 Gb. We were unable to get hardware requirements for VCF-kit as it failed to process the datasets and ran into out-of-memory errors.

When Prometheus was activated, CATE was configured to use a maximum of 40 CPU cores, and process 1,000 query regions at a time with only 100,000 SNPs being submitted to the GPU at once. CATE was recorded to have used 100% of the CPU with RAM usage between 30.73 Gb and 50 Gb. CATE on Prometheus was recorded to have a maximum of 70% of GPU usage with about 400 Mb of memory being used at maximum. There is a trade-off between Prometheus’ speed and its demand on resources. However, it should be noted that Prometheus’s usage of RAM can be limited by limiting the number of query regions it will have to process at a time and CATE’s Prometheus has the advantage of being configured based on resource availability.

### Neutrality test analysis of SARS-CoV-2 genome

In our analysis of the Delta variant from the GISAID SARS-CoV-2 database (**Supplementary materials 4.3**) we found that CATE’s results (**Supplementary materials 5.2.4**) coincided with that of existing literature. We observed that under the Tajima’s D test, the gene regions of the envelope protein, membrane protein, and Open Reading Frame 7b had less negative Tajima’s D values. These results coincide with the work conducted by Farkas et al. in 2021 (40). Additionally, consistent with existing work the neutrality tests showed that the gene regions of Open Reading Frame 1ab, Spike protein, and Nucleocapsid had an abundance of rare alleles due to the rise of mutations and selection sweeps in these regions during the pandemic (41, 42).

## DISCUSSION

Through this work, our goal was to build a novel computational framework for big data analysis of genome-wide genetic variation data, with potential applications beyond tests of neutrality. Our software CATE implements this framework proving its viability. CATE is scalable, resource-efficient, and fast. It has been built with the aim of reducing compute time through large-scale parallel processing of data. We achieve this level of parallel processing, using CUDA enabled NVIDIA GPUs, the CPU, and where available the SSD.

Large scale data collection leads to increased dimensionality, which leads to complications in the efficient use of computational resources when processing these expansive data sets. To address this a number of parallel processing solutions have been brought forward (17, 43). These include dedicated software built around multi-core processing as well as the expansion of statistical tools such as R to enable support for parallel processing. Most of these solutions limit themselves to the CPU, which at best contains tens of cores and hundreds of threads (43, 44). Modern GPUs on the other hand house over thousands to tens of thousands of cores on a single chip. This paradigm shift in parallel processing technology has resulted in the emergence of genomic software that show marked improvements in processing speeds using GPU based algorithms (24, 26, 27, 45).

However, the ability to process data is limited by the time elapsed for its access and retrieval. In an attempt to overcome this predicament, some modern solutions use indexing tools to navigate large files while others attempt to load sections of the repository into RAM (46). Despite this, to access the target region, the software has to sequentially navigate the repository, skipping over the data till the target region is reached (10, 46). With larger repositories loading the data into RAM becomes resource intensive. Our unique segmented file structure with our Compound Interpolated Search (CIS) algorithm addresses these obstacles while providing true random access to the data store. Due to the segmented nature of the data store, it enables the use of SSD technologies to access multiple data points simultaneously.

Our unified implementation of large-scale parallel processing using the GPU and CPU with our bespoke algorithms along with the augmentation of the data retrieval process has resulted in a framework when applied to our software CATE allows it to become magnitudes faster while applying little stress on hardware resources. At optimum conditions, CATE has proven to be on average over 180 times faster than conventional tools. This framework can be expanded beyond CATE, as a standard for genomic data storage and processing to be used in future software solutions.

CATE’s limitation lies in its reliance on CUDA enabled GPUs, a technology proprietary to the NVIDIA cooperation (24). However, these GPUs have steadily become a mainstream technology that is a standard in modern high-performance computing and is found in most personal computers. At present, CATE has 16 functions. We hope to further enhance these capabilities by adding more functionality and expand its support for different file formats. CATE is a command line tool and due to its compatibility with personal computing hardware, we hope to implement a GUI-based solution in the future.

The segmentation of variant call data sets in an organizable manner together with the use of parallel processing technologies such as the GPU is our novel proposition to solve the dilemmas of large-scale genomic data analysis. Through our software CATE we have proven this framework works as intended. CATE’s efficiency enables it to not only be fast but allows the processing of large-scale genomic data that would otherwise require the use of specialised computational systems on general-purpose computer hardware. Through these innovations, we hope to alleviate the strain of processing large-scale datasets and make CATE a standard tool in a bioinformatician’s arsenal.

## DATA AVAILABILITY

CATE is an open source and free software available for download on GitHub at: https://github.com/theLongLab/CATE

## ACKNOWLEDGEMENTS

Conceived the project: D.P. and Q.L. Designed the algorithm: D.P. Implementation and benchmarking: D.P., E.R., S.H. and E.H. Write the manuscript: D.P., Q.L., and C.D.H. with contribution from all the authors. Supervision: Q.L. and C.D.H.

