## Supplementary Materials for "CATE: A fast and scalable CUDA implementation to conduct highly parallelized evolutionary tests on large scale genomic data"

**LIST OF SUPPLEMENTARY FIGURES**

**Figure S - 1:** Detailed sequence diagram showing the life cycle of execution of evolutionary tests. It requires the user's input during the configuration of the parameter file and query regions. The program will then conduct the processing of the function and provide the user with a summarised output file. As shown CATE will provide the user with a continuous stream of notifications through the Command Line Interface (CLI) detailing its progress. ........ 5

**Figure S - 2.** The "IF" function has an additional three conditional statements added to it. As depicted in the figure this enables the algorithm to move away from the traditional equals conditional statement and capture ranges. Using the three additional statements linked through "OR" functions we can account for all possibilities. ........ 14

**Figure S - 3.** A sample snippet of code showing the conversion of the collected SNPs from a string vector into a char array during the Tajima's D calculation. The vector string of SNPs is concatenated end to end while keeping track of the start and stop position of each SNP. The concatenated string is then converted to a char array which can then be submitted to the GPU for parallel processing. Since we know the start and stop position of each SNP in the array, we can provide each thread with the relevant coordinates to process. ........ 16

**Figure S - 4:** Detailed view of the architecture of CATE's high-performance mode Prometheus. The grey dotted arrows signify when a dataset has been carried forward to a process and solid arrows indicate the flow of the command execution. As shown, it utilizes the GPU and three separate CPU-based multithreading blocks to conduct parallel processing. Multithreading block 1 is used to determine which segments are needed from the file hierarchy to satisfy the entire batch of query segments being processed. Multithreading block 2 filters and indexes the SNPs that have been processed by the current GPU round. Multithreading block 3 processes the evolutionary test for each query region in parallel. Prometheus can shift through multiple file reads to sequential reads based on the availability of SSD technologies The algorithm is designed to ensure that all these parallel processing techniques work together cohesively. ........ 19

**Table S - 30.** Detailed overview of the 11 codons that align with the outgroup regions and the population samples. Bases that have been subject to substitutions in the reference sequence are indicated in green. The SNPs that are fixed substitutions are demarcated in black for nonsynonymous mutations ($D_n$) and maroon for synonymous substitutions ($D_s$). The within species polymorphisms are denoted by grey and orange colours for nonsynonymous ($P_n$) and synonymous substitutions ($P_s$) respectively. As shown the substitution count is calculated by a parsimonious method, meaning they are all direct single base mutations. .......... 54
**Table S - 31.** CATE's output for the MK test. Comparison of the values with that of our example show that the algorithm works as intended. .......... 54

### SECTION 1. SOFTWARE OVERVIEW

CATE or CUDA Accelerated Testing of Evolution is a fast and scalable CUDA implementation to conduct highly parallelized evolutionary tests on large-scale genomic data.

CATE is available at the GitHub repository: https://github.com/theLongLab/CATE

#### 1.1. User interaction

Users will interact with CATE via the command line. They will enter the function they require with CATE's proprietary properties file and in certain instances where needed a gene file containing the coordinates of the query regions to be analyzed.

A detailed overview of the lifecycle of CATE is depicted through a sequence diagram in **Figure S - 1**. It shows the required prerequisites that need to be prepared before the execution of CATE as well as when these prerequisites will be used. CATE provides the user with a series of messages notifying its progress in the execution of the user-defined query regions as well as providing a summarised output file complete with the results from the test statistic.

A detailed breakdown of the steps required for CATEs execution:

1. Selection of the evolutionary test function to be executed.
2. Selection of the calculation mode (Window/ Gene mode).
3. If any other external files need to be prepared such as a gene file, they must be prepared. This is dependent on the function being executed.
4. Preparation of the parameters.json file.
5. Execution of CATE via the Command Line Interface (CLI).
    a) The user will pass the argument for the function and,
    b) The location of the parameters.json file.
6. While CATE is running it will provide the user with a series of messages updating the user of its progress in real-time.
7. Finally, the user will be provided with a tab-delaminated file of the results.

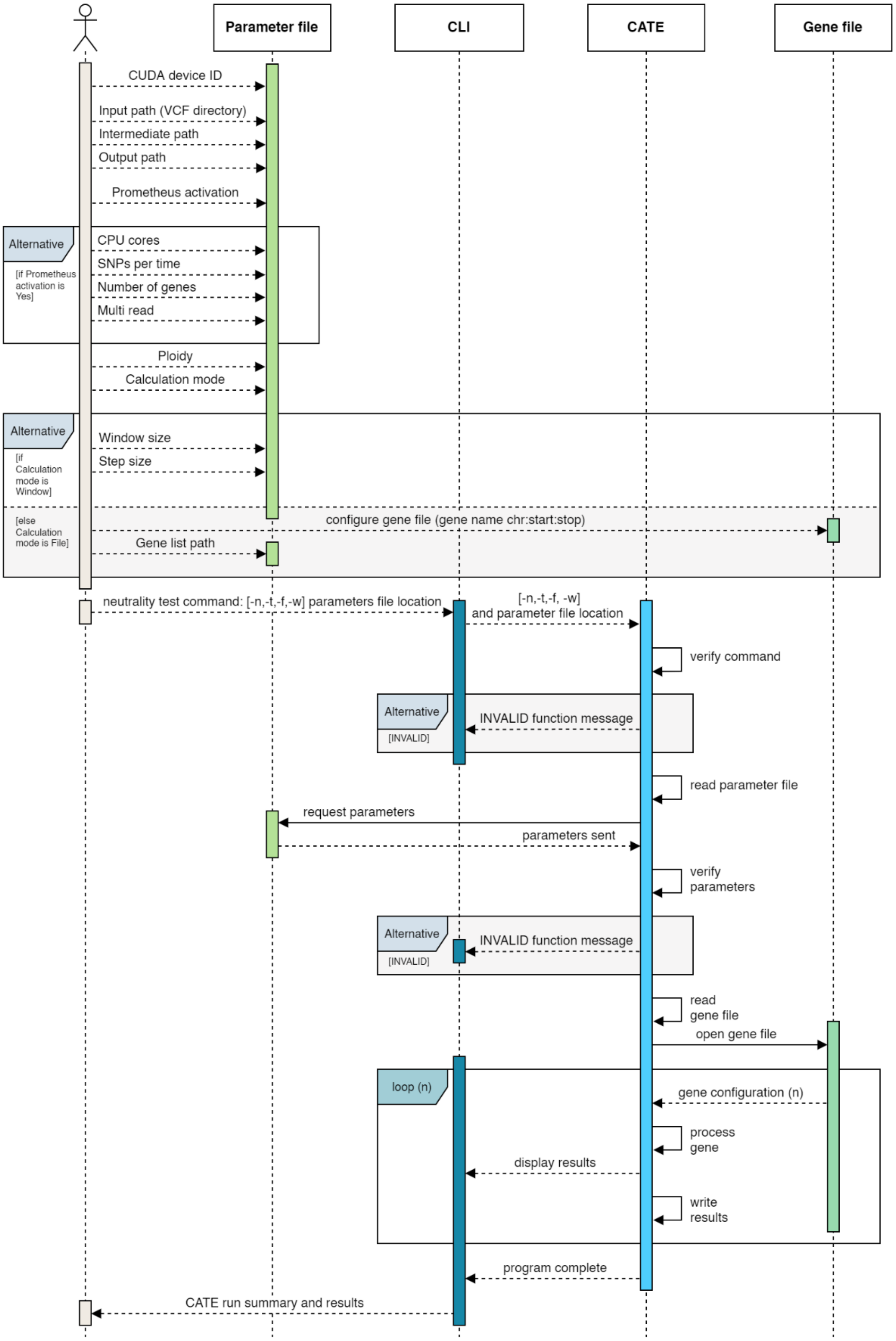

**Figure S - 1:** Detailed sequence diagram showing the life cycle of execution of evolutionary tests. It requires the user's input during the configuration of the parameter file and query regions. The program will then conduct the processing of the function and provide the user with a summarised output file. As shown CATE will provide the user with a continuous stream of notifications through the Command Line Interface (CLI) detailing its progress.

#### 1.2. Available functions

CATE has a total of 16 functions. Seven complementary tools designed for genomic data processing, and augmentation, seven functions dedicated for six different evolutionary tests, and two helper functions. A detailed description of these functions and their arguments are listed in **Table S - 1** below.

**Table S - 1.** A brief description of all available functions in CATE complete with the relevant command line arguments required to execute each function.

| Function name | Argument | | Description |
|---|---|---|---|
| ***Tools*** | | | |
| VCF splitter | -svcf | --splitvcf | Splits the parent VCF file into CATE's indexed format. Requires a population file to separate the VCF by populations. |
| FASTA splitter | -sfasta | --splitfasta | Can be used to extract a user specific single FASTA sequence from a merged FASTA file or to extract all sequences separately. |
| FASTA merger | -mfasta | --mergefasta | Merges multiple FASTA files in a folder into one singular FASTA file. |
| Gene extractor | -egenes | --extractgenes | Used to extract the FASTA sequences of target regions specified though CATE's gene file. Requires a reference genome in FASTA format. |
| GFF to Gene file converter | -g2g | --gff2gene | Used to extract the gene regions from a GFF file. It organises them into CATE's gene file format. |
| MAP to Gene file converter | -m2g | -- map2gene | Used to extract the SNP regions from a MAP file. It organises them into CATE's gene file format. |
| Haplotype extractor | -hapext | --hapfromvcf | Identifies unique haplotypes for different regions and reconstructs the FASTA sequences for each haplotype. Can be used to reconstruct the entire sequence population as well. |
| Print sample parameter file | -ppram | --printparam | Automatically generates a sample parameter file for the user. |
| ***Evolutionary tests*** | | | |
| Tajima's D statistics test | -t | --tajima | Calculates the Tajima's D statistic (1989). |
| Fu and Li statistics test | -f | --fuli | Calculates the Fu and Li's D, D*, F and F* statistics (1993). The F* statistic's vf* and uf* are calculated |

| | | | |
|---|---|---|---|
| | | | based on the corrected equations in Simonsen et al (1995). |
| Fay and Wu statistics test | -w | --faywu | Calculates the Fay and Wu's normalized H and E statistics (2006). |
| Neutrality tests | -n | --neutrality | Calculates the above three Neutrality tests (Tajima's Fu and Li's and Fay and Wu's) at once. |
| McDonald–Kreitman neutrality index | -m | --mk | Calculates the McDonald–Kreitman Neutrality Index (NI) (1991). |
| Fixation index | -x | --fst | Calculates the Fixation Index (Fst) (1965). |
| Extended Haplotype Homozygosity | -e | --ehh | Calculates the Extended Haplotype Homozygosity (EHH) (2002). |
| ***Helper functions*** | | | |
| CUDA device list | -c | --cuda | Lists all available CUDA devices. User can then use the CUDA ID to configure CATE. |
| Help menu | -h | --help | Prints the general help menu built into CATE. |

Each argument listed above has two options, an abbreviated and a long version. The arguments are not case sensitive. The abbreviated version follows a single hyphen (-) and the long version follows double hyphens (--).

#### 1.3. Parameter file

The user will configure CATE's function via a proprietary parameters file present in JSON format. The file is simple enough to be created by the user, and there is a built-in function in CATE (CLI argument is -ppram or --printparam) to create a sample parameter file to be edited by the user.

The list of all available parameters used to configure CATE are described in **Table S - 2** below. These parameters are case sensitive.

**Table S - 2.** Detailed descriptions of available parameters in CATE, their use and which functions they belong to.

| Parameter | Description |
|---|---|
| ***CUDA device*** | |
| CUDA Device ID | The CUDA device ID is used to configure CATE to which CUDA enabled GPU present in the system that it can use. |
| ***Directories*** | |
| Input path | The location of CATE's file hierarchy system. |
| Output path | The location to which all outputs will be written to. |
| Intermediate path | The location to which intermediate files such as log files will be written to. |
| ***Prometheus settings*** | |
| Prometheus activate | Boolean function (YES/ NO) states whether CATE should use Prometheus' high-performance mode or not. |
| CPU cores | Configures the number of CPU cores available for Prometheus to use. |
| SNPs per time | Configures the number of SNPs the GPU will process at a time. |
| Number of genes | Configures the number of query regions CATE will process at a time. |
| Multi read | Boolean variable (YES/ NO). States whether to read multiple files simultaneously or not. If YES SSD technologies are required to be present. |
| ***VCF ploidy*** | |
| Ploidy | Defines number of sets of chromosomes per individual organism. |
| ***Protein coding information*** | |
| Start codon(s) | Configurations of DNA codons that code for starting amino acids. |

| | |
|---|---|
| Stop codon(s) | Configurations of DNA codons that code for stop amino acids. |
| Genetic code | The genetic code for DNA to amino acids. |
| ***Neutrality and Fst calculation mode*** | |
| Calculation mode | Boolean variable (FILE/ WINDOW). Specifies if the calculation mode for determining the query regions. Either the gene file mode or traditional window mode. |
| ***Window mode parameters*** | |
| Window size | Size of the window within which the calculation will be conducted. |
| Step size | Size of the steps taken before the next window is reached. |
| ***Universal gene list file location*** | |
| Universal gene list | If all tests can be required to use the same gene list file, its location can be specified here.<br>The keyword, “universal” can be used to point the tests gene list here. |
| ***VCF splitter parameters*** | |
| Split mode | Boolean variable (CHR/ CTSPLIT).<br>CHR mode will split the VCF file by chromosome as well as extract out the GT column. It can also be used to specify the number of the reference and alternate alleles.<br>CTSPLIT will create CATE file structure by splitting the VCF file into file segments. VCF should only have the GT column data. |
| CHR individual summary | Used by CHR mode. At the end of the program’s execution, it will provide the user with a ratio of each samples coverage of each chromosome present within the final vcf. |
| Split cores | Configures the number of CPU cores available for the split function to use. |
| Split SNPs per_time_CPU | Configures the number of SNPs the CPU will process at a time. |
| Split SNPs per_time_GPU | Configures the number of SNPs the GPU will process at a time. |
| Population file path | The location of the tab delaminated file contained the samples population organisation. |
| Sample_ID Column number | Points to the column number (nonzero digit) which contains the sample ID that matches that of the VCF file. |

| | |
|---|---|
| Population_ID Column number | Points to the column number (nonzero digit) which contains the population ID. |
| Reference allele count | Number of alleles that can be present in the VCF’s reference column. |
| Alternate allele count | Number of alleles that can be present in the VCF’s alternate column. |
| SNP count per file | Maximum number of SNPs that will be in each segment. |
| MAF frequency | Defines the value of the Minor Allele Frequency (MAF), that will be used for the filtration of SNPs. |
| Frequency logic | Can be either one of five comparison operators.<br>= Equal<br>> Greater than<br>< Less than<br>>= Greater than or equal<br><= Less than or equal |
| ***FASTA splitter parameters*** | |
| Sequence | Contains the sequence ID of the sequence that needs to separate. If the keyword “ALL” is entered, then the entire file is split into separate sequence FASTA files. |
| Raw FASTA file | The original FASTA file containing all the sequences. |
| ***FASTA merger parameter*** | |
| FASTA files folder | The location of the folder that contains the separate FASTA files. |
| Merge FASTA path | The location to which the merged FASTA file should be written to. |
| ***Gene extractor*** | |
| Reference genome ex | The location of the FASTA format reference genome. |
| Extract gene list | The gene file location containing the list of query regions and their coordinates. Can use the keyword “universal” to point to the “Universal gene list” parameter. |
| ***GFF to Gene file converter*** | |
| GFF file | The location of the GFF file from which the gene’s coordinates will be extracted. |
| ***MAP to Gene file converter*** | |
| MAP file | The location of the MAP file from which the SNP coordinates will be extracted. |

| | |
|---|---|
| SNP prefix | Prefix that will be added in front of the SNP number, defining the SNP name. |
| ***Haplotype extractor*** | |
| Reference genome hap | The location of the FASTA format reference genome. |
| Population out | Boolean variable (YES/ NO). If YES, then the entire population's FASTA sequences will be generated. |
| Hap extract gene list | The gene file location containing the list of query regions and their coordinates. Can use the keyword "universal" to point to the "Universal gene list" parameter. |
| ***Neutrality tests parameter*** | |
| Neutrality gene list | The gene file location containing the list of query regions and their coordinates. Can use the keyword "universal" to point to the "Universal gene list" parameter. |
| ***Tajima's D parameters*** | |
| Tajima gene list | The gene file location containing the list of query regions and their coordinates. Can use the keyword "universal" to point to the "Universal gene list" parameter. |
| ***Fu and Li parameters*** | |
| Fu and Li gene list | The gene file location containing the list of query regions and their coordinates. Can use the keyword "universal" to point to the "Universal gene list" parameter. |
| ***Fay and Wu parameters*** | |
| Fay and Wu gene list | The gene file location containing the list of query regions and their coordinates. Can use the keyword "universal" to point to the "Universal gene list" parameter. |
| ***McDonald–Kreitman parameters*** | |
| Alignment mode | Boolean variable (GENE/ CHROM). Specifies whether the conducted alignment of the reference sequence with the outgroup was a Gene-Gene alignment or a Chromosome-Chromosome wide alignment. |
| ORF known | Boolean variable (YES/ NO). Specifies if CATE should automatically find the best OR in the specified region or that the user has already specified the ORF's coordinates. |
| Reference genome mk | The location of the FASTA format reference genome. |
| Alignment file | The location of the alignment file. |
| McDonald–Kreitman gene list | The gene file location containing the list of query regions and their coordinates. Can use the keyword "universal" to point to the "Universal gene list" parameter. |

| | |
|---|---|
| ***Fixation Index*** | |
| Population index file path | The location of the tab delaminated file, which contains the samples population organisation. |
| Fst gene list | The gene file location containing the list of query regions and their coordinates. Can use the keyword “universal” to point to the “Universal gene list” parameter. |
| Population ID | The list of populations that will be used to conduct the population wide global Fixation index calculation. |
| ***Extended Haplotype Homozygosity*** | |
| Range mode | Two main modes. The first enables the user to specify the core haplotype as a region (defined by a start and stop). The second enables the user to specify it as a SNP (a single position), so the core haplotypes will be 0 (reference allele) and 1 (alternate allele).<br>FILE/ FIXED modes. Specifies whether the extended haplotype region is a fixed displacement from the core site or has to separately determined using the gene file.<br>SNP/ BP modes. Here the core haplotype is a single SNP position. The extended haplotype region is specified in the number of SNPs or BPs that will be displaced in front of and behind the SNP position. The EHH values will then be calculated for the core haplotypes of 0 (reference allele) and 1 (alternate) for every allele in the extended region making up the extended haplotypes. |
| EHH FILE path | The gene file location (for FILE mode) containing the list of query regions and their coordinates. Can use the keyword “universal” to point to the “Universal gene list” parameter. |
| FIXED mode | Fixed displacement (for FIXED mode) of the core haplotype region to identify the extended haplotype region. |
| SNP default count | Used by the SNP mode. It determines the number of SNPs that will be located before and after the central SNP (core haplotype), thereby defining the extended region. |
| SNP BP displacement | Used by BP mode. It determines the number of base pairs that will be located before and after the central SNP (core haplotype), thereby defining the extended region. |

### SECTION 2. NOVEL IMPLEMENTATIONS

CATE attempts to solve the problem of latency in conducting evolutionary tests on expansive genomic data through the process of large-scale parallelization. To achieve this goal, we have decided to use the parallel processing capabilities of the GPU together with the CPU. Modern GPU architecture is specifically designed for large-scale parallelization. A typical GPU comes equipped with thousands of computing cores, well above their CPU counterparts (1). We have programmed the GPU using the CUDA (Compute Unified Device Architecture) framework developed by the NVIDIA corporation (2). CATE is also programmed to use multiprocessing technologies of the CPU, and in certain instances, it is extended to enable SSD technologies for parallelization in file reads.

Through this section, we aim to provide a more detailed breakdown of our algorithm to provide the user with a more comprehensive understanding of how we arrived at our goal.

#### 2.1. File hierarchy

The Variant Call File (VCF) has been designed specifically to store a population's genetic polymorphism (3). The file system, though designed to capture polymorphisms of large-scale datasets such as the 1000 Genome project, can become impractical to access especially when the number of variants stored become large. For instance, the 1000 Genome project has over 88 million variants spread across 23 chromosomes (4). Since programs and programming languages are designed to sequentially read through these files, it becomes a tedious process to manage multiple regions of interest. Most solutions to this problem are usually limited by the computer's RAM as they rely on loading large sections of the files to the onboard memory (5). Therefore, when designing a faster computational solution that is capable of working on minimal resources, this is the first obstacle that must be overcome.

Therefore, we have created our own file system for CATE. This file hierarchical system breaks the VCF file into segments and allows the indexing and sorting of the variant data by their position in the genome. It was also ensured that this file system does not deviate too greatly from the standard VCF files and if needed the original VCFs can be reconstructed. We have a built-in tool (VCF splitter) to enable the automatic creation of the file system. The file system was designed so that it can be created by a user with minimal supervision even in the absence of our inbuilt tool.

#### 2.2. Detailed overview of Compound Interpolated Search (CIS)

The Compound Interpolated Search (CIS) algorithm (**main text Figure 2**) is a bespoke algorithm designed specifically for CATE. The algorithm combines two traditional search algorithms: interpolation search and sequential search. Additionally, the structure of the algorithm enables the use of the CPU's multiprocessing capabilities. We use the interpolation search to capture a latch point which will then spawn two sequential searches to capture the rest of the segments that satisfy the needs of our region of interest.

As stated in the main text, certain components of the two search algorithms have been modified to accommodate CATE’s file hierarchy. CATE’s file hierarchy requires a single large VCF file to be broken down into segments. These segments have a fixed number of SNPs which ensures that the time taken to read through each one remains fairly constant. The segments are then labeled by the coordinates of the SNPs that they encompass. Simply said each segment's name will specify the lowest and highest position of the SNPs it contains. These segments are mutually exclusive meaning that they do not have overlaps with each other.

When a user enters a query region, they specify a range within which we have to look for polymorphic data. Therefore, unlike in a traditional search algorithm, we are not looking for a specific value within a dataset of singular values. Instead, we are trying to capture the segments that contain data that will satisfy our query range. Simply put, we are looking for ranges that will contain values within our query range. To achieve this purpose, we have modified the "IF" statements of both the interpolation search and the sequential search algorithms. **Figure S - 2** below depicts a graphical representation of the modified statement. Essentially by using a series of query regions with the "IF" function we are able to capture all possibilities of segments that will satisfy our query range.

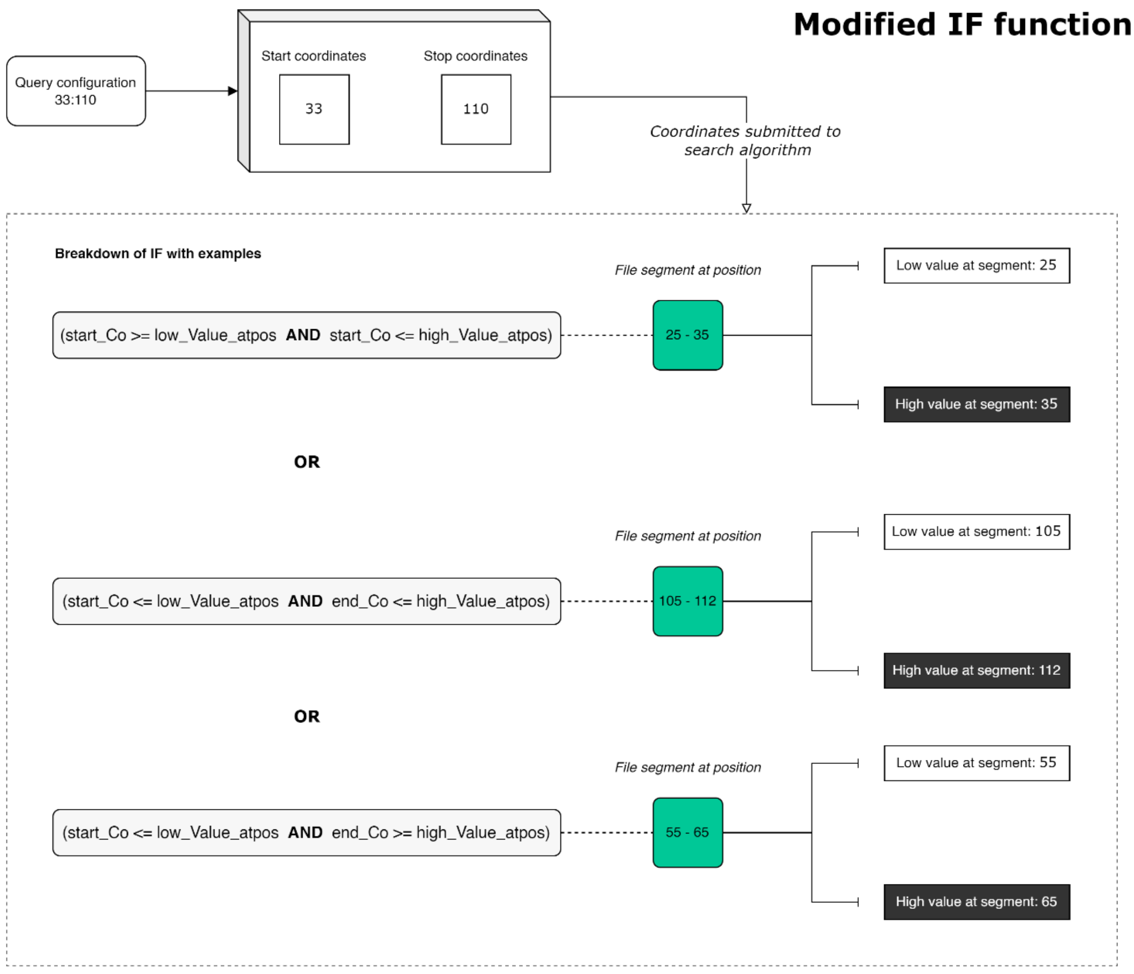

**Figure S - 2.** The "IF" function has an additional three conditional statements added to it. As depicted in the figure this enables the algorithm to move away from the traditional equals conditional statement and capture ranges. Using the three additional statements linked through "OR" functions we can account for all possibilities.

In addition to the modification of the “IF” function, we have also modified the position algorithm present in the interpolation search algorithm. The interpolation search algorithm has an added advantage over the binary search algorithm as it can weight your search value given that you know the highest and lowest values of your search space. Since CATE utilizes an indexed file hierarchy that is sorted by position these values are known. However, the traditional position algorithm is designed to be weighted by a single value of interest and when considering query regions, we have a range of interest between the start and end coordinates. Therefore, this positional equation was modified to be weighted by the start coordinate of the query region.

Capturing a single segment will not provide us with the complete set of polymorphic data required to process our query region. This is why once a latch point is found using the interpolation search algorithm, the CIS algorithm uses two sequential search algorithms to look for the remaining segments around this point.

#### 2.3. Transfer of data strings into the GPU

CUDA in its raw form cannot directly read strings of data; they need to be converted into character arrays. Therefore, we have come up with a solution to overcome this drawback and at the same time use it to better speed up the procedure of SNP processing in the GPU. A snippet of the code designed for this function is shown in **Figure S - 3**.

string to 1D char

```
/**
     * Administrative function responsible for collection of data required for Tajima's D.
     * 1. Conversion of SNP strings into char pointers for GPU accessability.
     * 2. Call GPU for extracting MAs (Minor allele) and MAF's (Minor Allele Frequencies).
     **/

    cout << "System is processing and filtering segregating site(s)" << endl;
    /**
     * @param num_segregrating_Sites is used to track the number of SNPs collected for the query region.
     * This track is vital for navigating through the data in the GPU. For the data is stored in the form of a 1D array.
     **/
    int num_segregrating_Sites = total_Segregrating_sites.size();

    /**
     * @param Seg_sites is used to stitch the SNP data end to end, before converting it to a char array.
     **/
    string Seg_sites = "";
    /**
     * @param site_Index is used to keep track of the start and ends of each SNP's data.
     **/
    int site_Index[num_segregrating_Sites + 1];
    site_Index[0] = 0;

    /**
     * Conversion of vector SNP information into a 1D array by concatenating the vector data into a single string.
     **/
    for (size_t i = 0; i < num_segregrating_Sites; i++)
    {
        Seg_sites.append(total_Segregrating_sites[i]);
        site_Index[i + 1] = site_Index[i] + total_Segregrating_sites[i].size();
    }

    /**
     * Final assignment of concatented string into a 1D char array.
     **/
    char *full_Char;
    full_Char = (char *)malloc((Seg_sites.size() + 1) * sizeof(char));
    strcpy(full_Char, Seg_sites.c_str());

    /**
     * RAM is released to prevent redundancy.
     **/
    total_Segregrating_sites.clear();

    // cuda_process_Seg(char *sites, int *index, int tot_Segregrating_sites, int *VALID_or_NOT, int *MA_count)
    /**
     * @param cuda_full_Char is used by the GPU. Is a COPY of full_Char.
     * @param cuda_site_Index is used by the GPU. Is a COPY of site_Index.
     *
     * * These 4 variables work together in all instances.
     **/
    char *cuda_full_Char;
    cudaMallocManaged(&cuda_full_Char, (Seg_sites.size() + 1) * sizeof(char));
    int *cuda_site_Index;
    cudaMallocManaged(&cuda_site_Index, (num_segregrating_Sites + 1) * sizeof(int));
    /**
     * Transfer of data to the GPU.
     **/
    cudaMemcpy(cuda_full_Char, full_Char, (Seg_sites.size() + 1) * sizeof(char), cudaMemcpyHostToDevice);
    cudaMemcpy(cuda_site_Index, site_Index, (num_segregrating_Sites + 1) * sizeof(int), cudaMemcpyHostToDevice);
```

**Figure S - 3.** A sample snippet of code showing the conversion of the collected SNPs from a string vector into a char array during the Tajima's D calculation. The vector string of SNPs is concatenated end to end while keeping track of the start and stop position of each SNP. The concatenated string is then converted to a char array which can then be submitted to the GPU for parallel processing. Since we know the start and stop position of each SNP in the array, we can provide each thread with the relevant coordinates to process.

For each test, we use a separate administrative function to convert the strings of data that CATE has collected from the VCF into char arrays. This provides the opportunity to concatenate the SNP sites into a singular array. Once the start and stop positions of each SNP is recorded the processing of the data can be parallelized in the GPU.

In doing so, CATE is capable of processing each SNP in a separate thread. This allows large-scale parallelization of the SNP processing step. Alternatively, this step would take a long time if done sequentially or even in the CPU. This is mainly due to the 1000s of SNPs that can be present in a region, especially for regions that span across a few thousand base pairs.

#### 2.4. Prometheus: CATE’s high-performance mode

CATE’s high-performance mode is designed to expedite CATE’s normal architecture by processing multiple query regions in unison through additional three regions of CPU based multithreading and one optional SSD based multithreading technique.

It has been implemented for the neutrality tests of Tajima’s D, Fu and Li's D, D*, F and F*, Fay and Wu’s H and E as they treat each segregating site independently. A detailed overview of CATE’s Prometheus architecture is depicted in **Figure S - 4**.

Once the user executes CATE for neutrality tests, it will recognize the activation of Prometheus. CATE will begin by collecting the start and stop positions of multiple query regions based on the user-specified parameter called number of genes. This controls the number of query regions that will be processed at a time. It will then determine the unique list of file segments required to satisfy this collection of query regions using the CIS algorithm. We carry out this filtration because multiple query regions might have common file segments. This step helps ensure that each file segment is present only once thereby preventing redundancy of segment file reads.

After the files have been obtained, CATE will proceed to read the files in their entirety. This is because at this point CATE does not know which SNP positions are required and which are not as multiple query regions are being processed. In this instance, if the hardware is equipped with an SSD or flash storage it can conduct multiple reads of multiple file segments at the same time.

Once the SNPs are committed into the computer memory Prometheus conducts its processing using a different strategy than CATE’s normal mode. In the normal mode only the SNP positions that satisfy the query range will be committed to the memory, this is because only a single range is processed at a time. However, in Prometheus as multiple query regions are processed together all SNP positions present in the selected segment files are committed to memory. Therefore, the positions of these SNPs in the genome are extracted at the same time each SNPs data points are extracted. This step is done in the GPU. This is similar to the normal modes GPU processing of each SNP with the addition of position extraction. Due to the large number of SNPs that can be present and due to the limitations in GPU memory, Prometheus comes with a parameter called SNPs per time which can be configured by the user. This will control the number of SNPs the GPU will process at a time. If the number of SNPs exceeds the per-run limit, the SNPs will be broken down into batches and processed via multiple GPU runs.

Once all the SNPs are processed, they are sorted by their genomic position. This enables CATE to search these SNP's using a variation of our CIS algorithm, where we replace the interpolation search with a binary search algorithm. We replace the interpolation search with the binary search algorithm because interpolation search requires a uniformly distributed list to be more efficient than its counterpart. In our testing we found that in instances where the query ranges where sporadic the interpolation search would cause a loss in efficiency. Therefore, to satisfy all instances we replaced it with the binary search algorithm. Once the SNPs are sorted each query region is processed in parallel by CPU multithreading.

When moving from one query region to the next in Window mode Prometheus checks to see if two consecutive regions utilise the same file segments. If so, it bypasses the data collection step moving straight to the calculation of the tests. This optimization reduces the processing time that would otherwise be wasted in redundant data collection and processing.

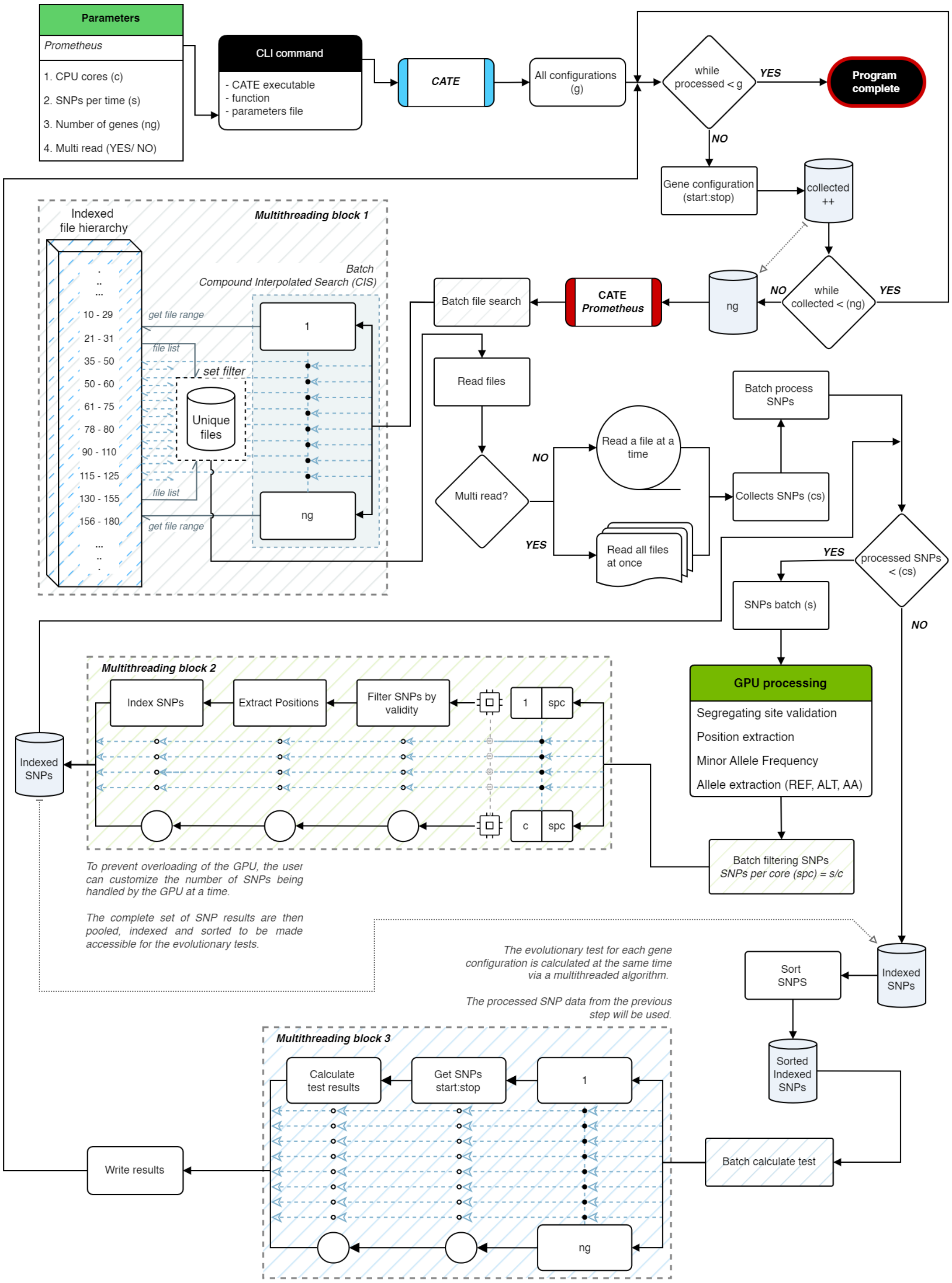

**Figure S - 4:** Detailed view of the architecture of CATE's high-performance mode Prometheus. The grey dotted arrows signify when a dataset has been carried forward to a process and solid arrows indicate the flow of the command execution. As shown, it utilizes the GPU and three separate CPU-based multithreading blocks to conduct parallel processing. Multithreading block 1 is used to determine which segments are needed from the file hierarchy to satisfy the entire batch of query segments being processed. Multithreading block 2 filters and indexes the SNPs that have been processed by the current GPU round. Multithreading block 3 processes the evolutionary test for each query region in parallel. Prometheus can shift through multiple file reads to sequential reads based on the availability of SSD technologies The algorithm is designed to ensure that all these parallel processing techniques work together cohesively.

#### 2.5. Resume capabilities

Large-scale data sets come with extended processing times and CATE is designed to work on personal computers as well as HPCs (High Performance Computer). Most HPC platforms come equipped with workload managers and schedulers such as SLURM (6). These schedulers may have restrictions on the time of execution, which means a particular software or execution can only run for a limited period before it will be terminated. To accommodate for this CATE comes equipped with an automatic resume function for all its evolutionary test functions. This enables CATE to detect a premature termination for a particular function and resume execution from the last successfully completed process. This feature is also useful in the event of an untimely hardware failure or an accidental program termination.

#### 2.6. Complete window or bin analysis

Due to CATE's file hierarchy, we overcome a drawback faced by most software such as PopGenome (5), vcf-kit (7), and vcftools (3) when conducting window or bin analysis. This drawback affects selscan (8) in its SNP-based analysis as well. Due to these software's inherent inability to handle large-scale datasets in their entirety, they resort to loading sections of the data for analysis.

Usually, they load regions of the chromosome at a time and conduct window-wise analyses on the loaded region. This data loading step would have to be done repeatedly if there are multiple regions that span across the different regions of the dataset, leading to delays in processing time.

CATE's file structure allows it to be aware of the first and last genomic positions of the chromosome whose variants have been called. This is because our naming convention for the file segments stores the positional range of the SNPs that are present within them. This allows CATE to overcome the drawback that plagues other software allowing complete windows of genetic regions to be analyzed without the user having to be aware of the dimensions of the region being analysed.

### SECTION 3. ALGORITHMIC IMPLEMENTATION OF EVOLUTIONARY TESTS

The implementation of evolutionary tests in computational code requires a thorough understanding of the mechanics of their statistical equations. This is further stressed when integrating it into the GPU. We have taken care to break down these equations into their constituents with minimal to no modifications or assumptions while isolating independent sections that can be parallelized. This approach has enabled us to generate CATE which preserves the integrity of these tests while dramatically improving their speed and resource efficiency in a computational setting.

In this section we will look at each of the six evolutionary tests and their implementations. Specifically, we will discuss any modifications or assumptions that have been done to these tests and their significance.

#### 3.1. Neutrality test statistics

In CATE we have implemented the complete solutions for three separate neutrality tests. These tests are used to identify if a region of interest is experiencing deviations from hypothesis of neutral or random evolution.

The three tests of neutrality are:

1. Tajima's D test statistic
2. Fu and Li's D, D*, F and F* test statistics
3. Fay and Wu's normalized H and E test statistics

CATE also has a final option which integrates all three of these tests to calculate them at the same time in a singular execution.

#### 3.1.1. Tajima's D test statistic

Tajima's test statistic (9) is used to measure if a genetic region is under selection or mutating randomly from one generation to the next. The strength of the statistic lies in the fact that it can be used to detect and even explain population size deviations from neutrality. Fumio Tajima formulated the statistic by taking into account the mean pairwise differences in a population and the Waterson's estimator. The Waterson's estimator is a measure of genetic diversity in a population. It divides the number of segregating sites by the time taken to for a particular site to become homozygous again by genetic drift or mutational loss alone ($S/a_1$) (10). According to Tajima if a populations genetic sequence is mutating neutrally without the effects of selection, these two statistics should be equal and therefore Tajima's D will be zero. This is because under neutrality both values are estimators for the mutation rate per base (11). Deviations from this value either in the positive or negative direction signify deviation.

The general equation for Tajima's D is shown below in ***equation a***:

$$D = \frac{\pi - {}^{S}/_{a_1}}{\sqrt{e_1 S + e_2 S(S-1)}} \; (\boldsymbol{equation\ a})$$

Where $\pi$ stands for the mean pairwise differences, $S$ is the number of segregating sites. $e_1$ and $e_2$ are prerequisites that are calculated from the sample size n. In the context of VCF's sample size is the total number of sequences in the population.

Before the determination of the number of mean pairwise differences and the number of segregating sites in a query region of interest Tajima's D requires the calculation of eight prerequisite values. The figure below shows these statistics and their implementations. We use a synergistic implementation of GPU and traditional C++ code to execute our aim.

To calculate the first two prerequisite values $a_1$ and $a_2$ (**Figure S - 5**) for Tajima's D the equations are solved using both the GPU and the CPU. Since both equations require the generation of a series of values that have to be summed together, the GPU is used to generate these series instantaneously through large-scale parallelization.

$$a_1 = \sum_{i=1}^{n-1} {}^{1}/_{i} \quad (equation\ 1)$$

$$a_2 = \sum_{i=1}^{n-1} {}^{1}/_{i^2} \quad (equation\ 2)$$

harmonic means

```
int N = N_tot - 1;
float *a1_CUDA, *a2_CUDA;
float *a1_partial, *a2_partial, a2;

a1_partial = (float *)malloc(N * sizeof(float));
a2_partial = (float *)malloc(N * sizeof(float));

cudaMallocManaged(&a1_CUDA, N * sizeof(int));
cudaMallocManaged(&a2_CUDA, N * sizeof(int));

a_Calculation<<<tot_Blocks, tot_ThreadsperBlock>>>(N, a1_CUDA, a2_CUDA);
  __global__ void a_Calculation(int N, float *a1_CUDA, float *a2_CUDA)
  {
      /**
       * Generate each value in the sequence for both a1 and a2
      **/
      int tid = threadIdx.x + blockIdx.x * blockDim.x;

      while (tid < N)
      {
        a1_CUDA[tid] = (float)1 / (tid + 1);
        a2_CUDA[tid] = (float)1 / ((tid + 1) * (tid + 1));
        tid += blockDim.x * gridDim.x;
      }
  }
 cudaDeviceSynchronize();

cudaMemcpy(a1_partial, a1_CUDA, N * sizeof(float), cudaMemcpyDeviceToHost);
cudaMemcpy(a2_partial, a2_CUDA, N * sizeof(float), cudaMemcpyDeviceToHost);

cudaFree(a1_CUDA);
cudaFree(a2_CUDA);

/**
  * Summation of the harmonic means
**/

a1 = 0;
a2 = 0;
for (size_t i = 0; i < N; i++)
{
  a1 += a1_partial[i];
  a2 += a2_partial[i];
}

cout << "a1\t: " << a1 << "\t"
     << "a2: " << a2 << endl;

free(a1_partial);
free(a2_partial);
```

**Figure S - 5.** For equation 1 and equation 2 CATE utilizes a combination of general CPU calculations with GPU-level arithmetic processing. The GPU is used to generate the series of values required for the final summation. Each thread in the GPU generates a single value in the series. This becomes highly useful in samples with large population sizes.

For the remaining six prerequisite values (**Figure S - 6**) we solve using CPU based implementations. They are general equations that do not require any form of parallelization. Due to their simplicity and interdependence (as the former value will be used in the latter equation) they are processed in a singular CPU thread.

$$b_1 = \frac{n + 1}{3(n - 1)} \qquad (equation\ 3)$$

$$b_2 = \frac{2(n^2 + n + 3)}{9n(n - 1)} \qquad (equation\ 4)$$

$$c_1 = b_1 - {}^{1}/_{a_1} \qquad (equation\ 5)$$

$$c_2 = b_2 - \frac{n + 2}{a_1 n} + \frac{a_2}{a_1^2} \qquad (equation\ 6)$$

$$e_1 = {}^{c_1}/_{a_1} \qquad (equation\ 7)$$

$$e_2 = {}^{c_2}/_{(a_1^2 + a_2)} \qquad (equation\ 8)$$

```
float b1 = (float)(N_tot + 1) / (3 * (N_tot - 1));
float b2 = (float)(2 * (pow(N_tot, 2.0) + N_tot + 3)) / ((9 * N_tot) * (N_tot - 1));

cout << "b1\t: " << b1 << "\t"
     << "b2: " << b2 << endl;

float c1 = b1 - (1 / a1);
float c2 = b2 - ((N_tot + 2) / (a1 * N_tot)) + (a2 / pow(a1, 2.0));

cout << "c1\t: " << c1 << "\t"
     << "c2: " << c2 << endl;

e1 = c1 / a1;
e2 = c2 / (pow(a1, 2.0) + a2);

cout << "e1\t: " << e1 << "\t"
     << "e2: " << e2 << endl;
```

**Figure S - 6.** Equations three to eight are CPU based simple C++ implementations. Unlike equations one and two the values are not independent and require the previous equation's result to be solved.

Data regarding the number of segregating sites ($S$) and the average pairwise differences ($\pi$) are calculated using the SNP collected using the VCF file hierarchy. The SNPs are processed in the GPU to determine the number of segregating sites and the pairwise differences are extracted using ***equation b*** which uses MAF. The MAF is also extracted during the GPU processing steps.

$$pairwise\ differences = MAF \times (1 - MAF) \times n^2\ (\boldsymbol{equation\ b})$$

To get the average pairwise differences (***equation d***) we divide the above value by the total number of possible pairwise query regions (***equation c***).

$$combinations\ = \frac{n!}{2!\,(n-2)!}\ (\boldsymbol{equation\ c})$$

$$average\ pairwise\ differences\ (\pi)\ = \frac{pairwise\ differences}{combinations}\ (\boldsymbol{equation\ d})$$

For the other tests, we use a similar approach. In instances where there are deviations from the standard or novelties in the implementation of code, we will discuss them in the remainder of this section.

#### 3.1.2. Fu and Li's D, D*, F and F* test statistics

The Fu and Li statistics similar to the Tajima's D statistic are measures of neutrality of selection. However, in contrast to Tajima's D, Fu, and Li by accounting for the age of mutations present in a population. To do so Fu and Li account for singleton mutations and outgroup mutations. This is a pretty straightforward calculation given the required polymorphic data (12).

When calculating the Fu and Li statistics we have made one assumption (stated below) in CATE as well as the adaptation of a correction done by Simonsen et al (1995) to the original equation of $v_{f*}$ appearing in the publication by Fu and Li (1993) (12, 13).

**Assumptions**

1. When assuming the outgroup allele, to account for singletons with an outgroup CATE will take the Ancestral Allele (AA) present for that SNP in the VCF file. The AA value is found in the INFO column of the VCF file.

2. In the absence of the AA information CATE will take the reference allele as the AA.

#### 3.1.3. Fay and Wu's normalized H and E statistics

Fay and Wu's test statistics are advancements of Tajima's D. Similar to Fu and Li's it makes uses of outgroup data. Likewise, we have carried the previous assumption (Section 3.1.2) associated with AA detection here as well. The strength of Fay and Wu statistics is that it helps identify genetic sequences under positive selection. The statistical implementations found in CATE are the 2005 normalised solution for Fay and Wu's original H statistic in 2000 and the normalised E statistic (14).

#### 3.2. Fixation Index ($F_{st}$)

Fixation index is a measure of differentiation between populations by a measure of genetic variance. It is used to measure the degree of speciation between populations. CATE has a full population wide implementation of the statistics. This allows the user to measure the fixation index between multiple populations. $F_{st}$ is calculated by measuring the difference between the expected heterozygosities in subpopulations and the expected heterozygosities in the overall population (15).

**Assumptions**

1) Characteristically, $F_{st}$ is calculated per segregating site. However, in events where there is the specification of a genetic region, which usually has more than one segregating site; CATE will provide the user with the average $F_{st}$ per segregating site.
   CATE will divide the sum total of all Fst values for each segregating site in the specified region, proceed to divide this total by the number of segregating sites and provide the user with an average value.

#### 3.3. Extended Haplotype Homozygosity (EHH)

Haplotype based tests such as EHH are being used to determine the forces of selection acting on a region of interest in relation to the age of the surrounding genetic structure. EHH was developed by Sabeti et al in 2002 (16). This long-range haplotype testing procedure is used to identify regions (core haplotype) that are recently facing effects of positive selection. EHH weighs the effects of recombination against a core haplotype and its surrounding extended haplotype region. Under the neutral theory of evolution, if only genetic drift is in effect then the surrounding linkage disequilibrium would be low due to higher levels of recombination around a core haplotype and an allele of interest would take a relatively long time to reach high frequencies in a population (17). However, under the effects of positive selection linkage disequilibrium is preserved due to lower recombination events in an effort to preserve a beneficial core haplotype mutation. This phenomenon is exploited by the EHH statistic to identify regions undergoing recent selection pressures (18).

In CATE we also provide the user with two options for calculating the EHH value. The first focusses on a single SNP as the core haplotype and measures the degradation of the EHH as you move the extended haplotype region further away and the second is the ability to define a core haplotype region which a specified extended region around which the EHH can be measured. In a simple sense, the algorithm of EHH is designed to identify the number of unique core haplotypes present as well as the probability that two randomly chosen chromosomes would be homozygous in the extended region for that core haplotype (16, 19). Each core haplotype will have a separate EHH value.

#### 3.4. McDonald–Kreitman test

The McDonald-Kreitman (MK) test introduced in 1991 measures the effects of evolution within and between species by accounting for the base substitution that occurs in the coding region of genes (20). A gene's coding region is comprised of an Open Reading Frame (ORF) that codes for the amino acid sequence of the ensuing protein.

An amino acid is the result of a translation of a trio of nucleotides called a codon and multiple codon variations can code for the same amino acid. The MK test looks at these codons for variations, both between species and within species. It then estimates which base substitutions would lead to changes or conservations in the translated amino acid. Synonymous substitutions that conserve the translated amino acids are considered neutral and those that lead to changes in the translated amino acid are called nonsynonymous substitutions. Changes within species are polymorphic substitutions and changes between species that were fixed within the species are fixed substitutions (21, 22).

As such the MK test accounts for four constraints as shown in **Table S - 3**. The MK test uses these parameters to formulate the Neutrality Index (NI) statistic, stated in ***equation e*** (20, 21).

**Table S - 3.** The MK test accounts for the following four factors. $D_s$ and $P_s$ are the number of synonymous substitutions between species and within species respectively per gene and $D_n$ and $P_n$ are the number of nonsynonymous substitutions between species and within species respectively per gene.

| | Fixed (between species) | Polymorphic (within species) |
|---|---|---|
| **Synonymous** | $D_s$ | $P_s$ |
| **Nonsynonymous** | $D_n$ | $P_n$ |

$$Neutrality\ Index\ (NI) = \frac{P_n/P_s}{D_n/D_s}\ (\boldsymbol{equation\ e})$$

The NI value can then be used to make deductions on neutrality and the effects of selection on a gene. NI values of one signify neutral selection. In such an instance all nonsynonymous mutations are considered to be neutral. If the NI value is greater then the gene is experiencing negative selection or an excess of polymorphisms within the species that do not spread across species into divergence. The excess of polymorphisms occurs in an attempt to remove the deleterious mutations that may already be present. A NI value of less than one would imply divergence by an excess of nonsynonymous mutations meaning positive selection (21).

**Assumptions**

1. In instances where the user is not affirmative of the ORF of the query region, they can use CATE to automatically predict a putative ORF. For this, we use the following three steps to identify a putative ORF region (23, 24).

    a. Locate base triads (codons) corresponding to start codons to determine the start points of potential ORFs. This is done by starting at the beginning of the query region and frameshifting by one base until the end of the gene sequence. With each shift, the presence of a start codon at the beginning of the frame is checked. By this method, we are able to find all available start codons on the sequence.

    b. For each of these start codon points, the frame is read in triads from that point till a stop codon is reached. The length of the start codon to stop codon in each frame is recorded.

    c. Finally, the frame with the longest sequence is selected as it has been proven that the longest sequence has the highest likelihood of being the ORF.

2. A VCF captures the polymorphic sites in a population that deviate from a reference genome. Therefore, when processing the MK test the consideration of these sites must be carefully accounted for. We assume that if a base position is present in the VCF and has a nonzero MAF it is a polymorphic site. If the base position is absent in the VCF or has a zero MAF it is assumed to be fixed in the population.

3. In mutations, either synonymous or nonsynonymous, the base substitution is considered to be parsimonious. Essentially, it will be considered as a single, direct base change.

### SECTION 4. DETAILS ON BENCHMARKING OF CATE

Benchmarking of CATE was performed to ensure its accuracy, speed and resource efficiency. CATE was benchmarked against three repositories across different test types. These databases were selected due to the unique test challenges that each one would provide.

The chosen repositories and the justifications for their selection are as follows:

#### 1. 1000 Genomes project

The 1000 Genomes project data repository is one of the most complete and expansive datasets currently available (4, 19). It will provide us with an extreme test case for CATE's resource efficiency and speed. We also used this dataset to evaluate CATE's accuracy against other software.

#### 2. 1001 Genomes project

The 1001 Genomes Project provides detailed whole-genome sequence variation in at least 1001 strains of the plant *Arabidopsis thaliana*. However, the limitations of the second-generation sequencing technologies that were used have resulted in massive numbers of short sequence fragments that must be aligned to a reference genome, which leads to missing large or complex structural variants and simple variants inside complex variants (25). This enabled us to test CATE in the presence of incomplete and non-ideal datasets.

#### 3. GISAID SARS-CoV-2 dataset

We selected the GISAID dataset's SARS-CoV-2 database to create a variant call dataset to analyse how CATE would handle large number sizes (over 9600 sequences were used) and different ploidy values. Both datasets of the 1000 Genomes project and 1001 Genomes project uses *Homo sapiens* and *Arabidopsis thaliana* which are diploid (ploidy is 2) organisms. SARS-CoV-2 is a haploid organism (ploidy is 1).

#### 4.1. 1000 Genomes Project

The 1000 Genomes project was used to test all three factors of accuracy, speed, and efficiency.

##### 4.1.1. Creating CATE's file structure

The creation of CATE's unique file structure is one of the crucial factors that enables its speed and resource efficiency. Therefore, to benchmark this function the VCFs of all 22 autosomal chromosomes from the 1000 Genomes were used. CATE's was tested using a MAF filter (set at MAF greater than 0.05) and without a filter. The details are detailed below in **Table S - 4** below.

**Table S - 4.** A detailed breakdown of the number of SNPs processed per chromosome.

| Chromosome | Number of SNPs |
|---|---|
| 1 | 6468095 |
| 2 | 7081601 |
| 3 | 5832277 |
| 4 | 5732586 |
| 5 | 5265764 |
| 6 | 5024120 |
| 7 | 4716716 |
| 8 | 4597106 |
| 9 | 3560688 |
| 10 | 3992220 |
| 11 | 4045629 |
| 12 | 3868429 |
| 13 | 2857917 |
| 14 | 2655068 |
| 15 | 2424690 |
| 16 | 2697950 |
| 17 | 2329289 |
| 18 | 2267186 |
| 19 | 1832507 |
| 20 | 1812842 |
| 21 | 1105539 |
| 22 | 1103548 |
| **Total** | **81271767** |
| **Average** | **3694171.23** |

The parameters used by CATE's vcf splitter for the analysis were as follows:

- **Split cores** **:** 40
- **Split SNPs per_time_CPU** **:** 500000
- **Split SNPs per_time_GPU** **:** 100000
- **SNP count per file** **:** 10000
- **Reference allele count** **:** 1
- **Alternate allele count** **:** 1

The above parameters were consistent for both with and without filter runs. In the with filter run, two additional parameters were set. They were as follows:

- **MAF frequency** **:** 0.05
- **Frequency logic** **:** >

The parameters were configured using CATE's parameter JSON file.

##### 4.1.2. Prometheus high performance mode

The following parameters were used by CATE for the testing of its high-performance mode Prometheus.

- **CPU cores** **:** 40
- **SNPs per time** **:** 1000
- **Number of genes** **:** 100000
- **Multi read** **:** Yes

#### 4.1.3. All genes test type

The genes for each chromosome were obtained via the R package biomaRt from Bioconductor (26).

**Table S - 5.** A detailed breakdown of the number of query regions processed per chromosome in all genes test type in the 1000 Genomes Project.

| Chromosome | Number of query regions |
|---|---|
| 1 | 5363 |
| 2 | 4047 |
| 3 | 3101 |
| 4 | 2563 |
| 5 | 2859 |
| 6 | 2905 |
| 7 | 2876 |
| 8 | 2386 |
| 9 | 2323 |
| 10 | 2260 |
| 11 | 3208 |
| 12 | 2818 |
| 13 | 1217 |
| 14 | 2244 |
| 15 | 2080 |
| 16 | 2343 |
| 17 | 2903 |
| 18 | 1127 |
| 19 | 2910 |
| 20 | 1317 |
| 21 | 736 |
| 22 | 1263 |
| **TOTAL** | **54849** |

#### 4.1.4. Window test type

For window wise analysis the step size and window size were both set to 10,000 base pairs (bp).

**Table S - 6.** A detailed breakdown of the number of query regions processed per chromosome in window test type with an equal step and window size of 10,000 in the 1000 Genomes Project.

| Chromosome | Number of query regions |
|---|---|
| 1 | 24924 |
| 2 | 24318 |
| 3 | 19791 |
| 4 | 19104 |
| 5 | 18090 |
| 6 | 17100 |
| 7 | 15912 |
| 8 | 14630 |
| 9 | 14114 |
| 10 | 13547 |
| 11 | 13488 |
| 12 | 13379 |
| 13 | 9609 |
| 14 | 8829 |
| 15 | 8253 |
| 16 | 9024 |
| 17 | 8120 |
| 18 | 7801 |
| 19 | 5906 |
| 20 | 6291 |
| 21 | 3871 |
| 22 | 3520 |
| **TOTAL** | **279621** |

##### 4.1.5. Sliding window test type

In sliding window analysis, the step size was 1 SNP, and the window size was 10,000 base pairs.

**Table S - 7.** A detailed breakdown of the number of query regions processed per chromosome in sliding window test type, with a window size of 10,000 in the 1000 Genomes Project.

| Chromosome | Number of query regions |
|---|---|
| 1 | 6176956 |
| 2 | 6764431 |
| 3 | 5567100 |
| 4 | 5462679 |
| 5 | 5022415 |
| 6 | 4784616 |
| 7 | 4503842 |
| 8 | 4404143 |
| 9 | 3405258 |
| 10 | 3812840 |
| 11 | 3865353 |
| 12 | 3685951 |
| 13 | 2719009 |
| 14 | 2530383 |
| 15 | 2313186 |
| 16 | 2588039 |
| 17 | 2220009 |
| 18 | 2164507 |
| 19 | 1747024 |
| 20 | 1733485 |
| 21 | 1049984 |
| 22 | 1051673 |
| **TOTAL** | **77572883** |

#### 4.2. 1001 Genomes Project

In the 1001 Genomes project, we analysed the variant call data of 5 chromosomes from *A. thaliana*.

##### 4.2.1. All genes test type

The genes of *A. thaliana* and the were obtained via The Arabidopsis Information Resource (TAIR) (27).

**Table S - 8.** A detailed breakdown of the number of query regions processed per chromosome in all genes test type in the 1001 Genomes Project.

| Chromosome | Number of query regions |
|---|---|
| 1 | 7508 |
| 2 | 4470 |
| 3 | 5650 |
| 4 | 4308 |
| 5 | 6559 |
| **TOTAL** | **28495** |

##### 4.2.2. Window test type

For window wise analysis the step size and window size were both set to 10,000 base pairs (bp).

**Table S - 9.** A detailed breakdown of the number of query regions processed per chromosome in window test type in the 1001 Genomes Project.

| Chromosome | Number of query regions |
|---|---|
| 1 | 3043 |
| 2 | 1964 |
| 3 | 2347 |
| 4 | 1860 |
| 5 | 2698 |
| **TOTAL** | **11912** |

##### 4.2.3. Sliding window test type

In sliding window analysis, the step size was 1 SNP, and the window size was 10,000 base pairs.

**Table S - 10.** A detailed breakdown of the number of query regions processed per chromosome in sliding window test type in the 1001 Genomes Project.

| Chromosome | Number of query regions |
|---|---|
| 1 | 37080 |
| 2 | 63828 |
| 3 | 189685 |
| 4 | 56473 |
| 5 | 37720 |
| **TOTAL** | **384786** |

#### 4.3. GISAID SARS-CoV-2 dataset

The creation of the SARS-CoV-2 variant call dataset was as follows.

A total of 9600 sequences of the Delta variant of SARS-CoV-2 collected between January 1$^{st}$, 2021, and December 31$^{st}$, 2021 obtained via Oxford Nanopore Technologies were downloaded through the GISAID database. Using the reference genome (Accession ID: NC_045512.2) of SARS-CoV-2 variant calling was conducted using the tool SARS-CoV-2-freebayes (28).

The three neutrality tests of Tajima's D, Fay and Wu test statistics and Fu and Li statistics were conducted on the resultant data. We conducted a comparative evaluation of the results obtained via CATE against existing work.

### SECTION 5. DETAILED OUTLINE OF RESULTS

#### 5.1. Creating CATE's file structure

When creating the file structure using the data from the 1000 Genomes project CATE averaged a per chromosome time of about 11.09 minutes in the absence of MAF filtration and about 6.34 minutes in the presence of the MAF filter (**Table S - 11**). The faster time in the presence of the filter is believed to be caused by the reduced number of SNPs that have to be written to the hard disk post processing.

**Table S - 11.** Total run time taken by CATE to create the file structure for each chromosome.

| Chromosome | Without MAF filter (s) | With MAF filter (s) |
|---|---|---|
| 1 | 1141 | 579 |
| 2 | 1251 | 641 |
| 3 | 1037 | 580 |
| 4 | 1006 | 597 |
| 5 | 903 | 548 |
| 6 | 921 | 480 |
| 7 | 825 | 538 |
| 8 | 854 | 458 |
| 9 | 604 | 455 |
| 10 | 737 | 429 |
| 11 | 714 | 455 |
| 12 | 734 | 359 |
| 13 | 527 | 278 |
| 14 | 473 | 309 |
| 15 | 422 | 229 |
| 16 | 547 | 288 |
| 17 | 428 | 235 |
| 18 | 436 | 272 |
| 19 | 357 | 185 |
| 20 | 319 | 190 |
| 21 | 196 | 134 |
| 22 | 212 | 126 |
| **Total runtime (s)** | **14644** | **8365** |
| **Average runtime (s)** | **665.64** | **380.23** |

#### 5.2. Neutrality tests

CATE was significantly faster than its competitors, in all tested scenarios. Detailed overviews of the tests are presented in the tables below.

##### 5.2.1. Validation of Tajima's D algorithm

Validation of the Tajima's D algorithm was conducted by the comparison of CATE's results against Pop Genome (5) and VCF-kit (7) using the Variant Call File for Chromosome 1 from the 1000 Genomes dataset. The results are shown in **Table S - 12**.

**Table S - 12:** Comparison of results for the Tajima's D test statistic of CATE against Pop Genome and VCF-kit.

| Coordinates | CATE | Pop Genome | VCF-kit |
|---|---|---|---|
| 10000:20000 | -1.77135 | -1.7713456 | -1.7713455644258664 |
| 20000:30000 | -1.26762 | -1.2676153 | -1.2676153011110054 |
| 30000:40000 | -0.82768 | -0.8276844 | -0.82768440289893 |
| 40000:50000 | -1.76828 | -1.7682825 | -1.768282534790013 |

##### 5.2.2. Validation of Fu and Li algorithm

Validation of the algorithms for Fu and Li's D and F was conducted by the comparison of CATE's results against Pop Genome using the Variant Call File for Chromosome 1 from the 1000 Genomes dataset. The results for Fu and Li's D and F are shown in **Table S - 13** and **Table S - 14** respectively.

**Table S - 13:** Comparison of results for the Fu and Li's D test statistic of CATE against Pop Genome.

| Coordinates | CATE | Pop Genome |
|---|---|---|
| 10000:20000 | -6.31802 | -6.2945046 |
| 20000:30000 | -0.89188 | -0.8915380 |
| 30000:40000 | -3.48428 | -3.4831865 |
| 40000:50000 | -1.29946 | -1.2976030 |

**Table S - 14:** Comparison of results for the Fu and Li's F test statistic of CATE against Pop Genome.

| Coordinates | CATE | Pop Genome |
|---|---|---|
| 10000:20000 | -4.65593 | -4.647592486 |
| 20000:30000 | -1.26663 | -1.266067945 |
| 30000:40000 | -3.1361 | -3.134510306 |
| 40000:50000 | -1.8793 | -1.877287041 |

#### 5.2.3. Neutrality tests run times

Using the 1000 Genomes dataset a test of all available functions and configurations on CATE was conducted in comparison with the existing tools PopGenome and VCF-kit. The results of these analyses are depicted in **Table S - 15**.

**Table S - 15.** Detailed overview of each test conducted on the 1000 Genomes dataset for the neutrality functions available on CATE against PopGenome, including the number of query regions (Qr.) that were analysed, the average time taken to process the entire length of an individual chromosome (Avg. chr.) and the total time taken to process the entire dataset.

| Software | Test type | Evolutionary test | Qr. | Avg. chr. (s) | Total time (s) |
|---|---|---|---|---|---|
| CATE normal mode | All genes | Neutrality full | 54,849 | 621.68 | 13,677 |
| | | Tajima's D | | 539.05 | 11,859 |
| | | Fay and Wu statistics | | 458.45 | 10,086 |
| | | Fu and Li statistics | | 525.86 | 11,569 |
| | Window | Neutrality full | 279,621 | 409.64 | 9,012 |
| | | Tajima's D | | 687.09 | 15,116 |
| | | Fay and Wu statistics | | 632.64 | 13,918 |
| | | Fu and Li statistics | | 404.00 | 8,888 |
| CATE Prometheus mode | All genes | Neutrality full | 54,849 | 78.18 | 1,720 |
| | | Tajima's D | | 79.14 | 1,741 |
| | | Fay and Wu statistics | | 78.68 | 1,731 |
| | | Fu and Li statistics | | 79.05 | 1,739 |
| | Window | Neutrality full | 279,621 | 70.32 | 1,547 |
| | | Tajima's D | | 72.82 | 1,602 |
| | | Fay and Wu statistics | | 72.86 | 1,603 |
| | | Fu and Li statistics | | 69.45 | 1,528 |
| | Sliding window | Neutrality full | 77,572,883 | 689.50 | 15,169 |
| | | Tajima's D | | 700.77 | 15,417 |
| | | Fay and Wu statistics | | 664.68 | 14,623 |
| | | Fu and Li statistics | | 676.05 | 14,873 |
| PopGenome | All genes | Neutrality full | 54,849 | 14,229.95 | 313,059 |
| | Window | Neutrality full | 279,621 | 3,617.41 | 79,583 |

The speed was also evaluated using the 1001 genomes dataset for all three test types as well under CATE's normal mode. The result of this analysis is depicted in **Table S - 16**.

**Table S - 16.** Detailed overview of each test conducted on the 1001 Genomes dataset for the neutrality functions available on CATE, including the number of query regions (Qr.) that were analysed, the average time taken to process the entire length of an individual chromosome (Avg. chr.) and the total time taken to process the entire dataset.

| Software | Test type | Evolutionary test | Qr. | Avg. chr. (s) | Total time (s) |
|---|---|---|---|---|---|
| CATE normal mode | All genes | Neutrality full | 28,495 | 40.00 | 200 |
| | | Tajima's D | | 38.60 | 193 |
| | | Fay and Wu statistics | | 37.20 | 186 |
| | | Fu and Li statistics | | 39.60 | 198 |
| | Window | Neutrality full | 11,912 | 19.20 | 96 |
| | | Tajima's D | | 19.20 | 96 |
| | | Fay and Wu statistics | | 20.60 | 103 |
| | | Fu and Li statistics | | 27.00 | 135 |
| | Sliding window | Neutrality full | 384,786 | 471.00 | 2,355 |
| | | Tajima's D | | 487.60 | 2,438 |
| | | Fay and Wu statistics | | 443.20 | 2,216 |
| | | Fu and Li statistics | | 442.20 | 2,211 |

#### 5.2.4. GISAID SARS-CoV-2 dataset

For the analysis of the SARS-CoV-2 data we analysed the 11 genes depicted in **Table S - 17**.

**Table S - 17.** Details of the selected 11 genes from the SARS-CoV-2 virus' genome.

| Gene ID | Gene name |
|---|---|
| GU280_gp01 | Open Reading Frame 1ab |
| GU280_gp02 | S protein or Spike protein. |
| GU280_gp03 | Open Reading Frame 3a |
| GU280_gp04 | E protein or Envelope protein |
| GU280_gp05 | M protein or Membrane protein |
| GU280_gp06 | Open Reading Frame 6 |
| GU280_gp07 | Open Reading Frame 7a |
| GU280_gp08 | Open Reading Frame 7b |
| GU280_gp09 | Open Reading Frame 8 |
| GU280_gp10 | N protein or Nucleocapsid |
| GU280_gp11 | Open Reading Frame 10 |

Following the neutrality test analysis, we conducted a comparative analysis with currently present literature to validate our results. The graphical representations of the results from CATE are shown in **Figure S - 7**.

As depicted in **Figure S - 7** (**A**) we observed that under the Tajima's D test the gene regions of GU280_gp04 (envelope protein), GU280_gp05 (membrane protein) and GU280_gp08 (Open Reading Frame 7b) had less negative Tajima's D values. These results coincide with the work conducted by Farkas *et al.* in 2021 (29). These results are also shown in the Fay and Wu values (**Figure S - 7** (**B**)). This suggests that in contrast to the rest of the genes they may be undergoing balancing selection.

From the Fu and Li test statistics as depicted in **Figure S - 7** (**C**) we observed three genes under significantly high negative Fu and Li values when compared to the surrounding genes. This could signify an abundance of rare alleles (12). They were GU280_gp01 (Open Reading Frame 1ab), GU280_gp02 (S protein or Spike protein) and GU280_gp10 (N protein or Nucleocapsid). The gene region of GU280_gp01 is composed of two Open Reading Frames (ORFs) respectively ORFa and ORFb. In the course of the COVID19 pandemic, it has been identified that this ORF1ab region has experienced the highest level of mutation in the SARS-CoV2 virus genome (30). This holds true for the Spike protein and Nucleocapsid phosphoproteins (31, 32). Therefore, these genes can have an abundance of rare allele mutations hence the negative Fu and Li values.

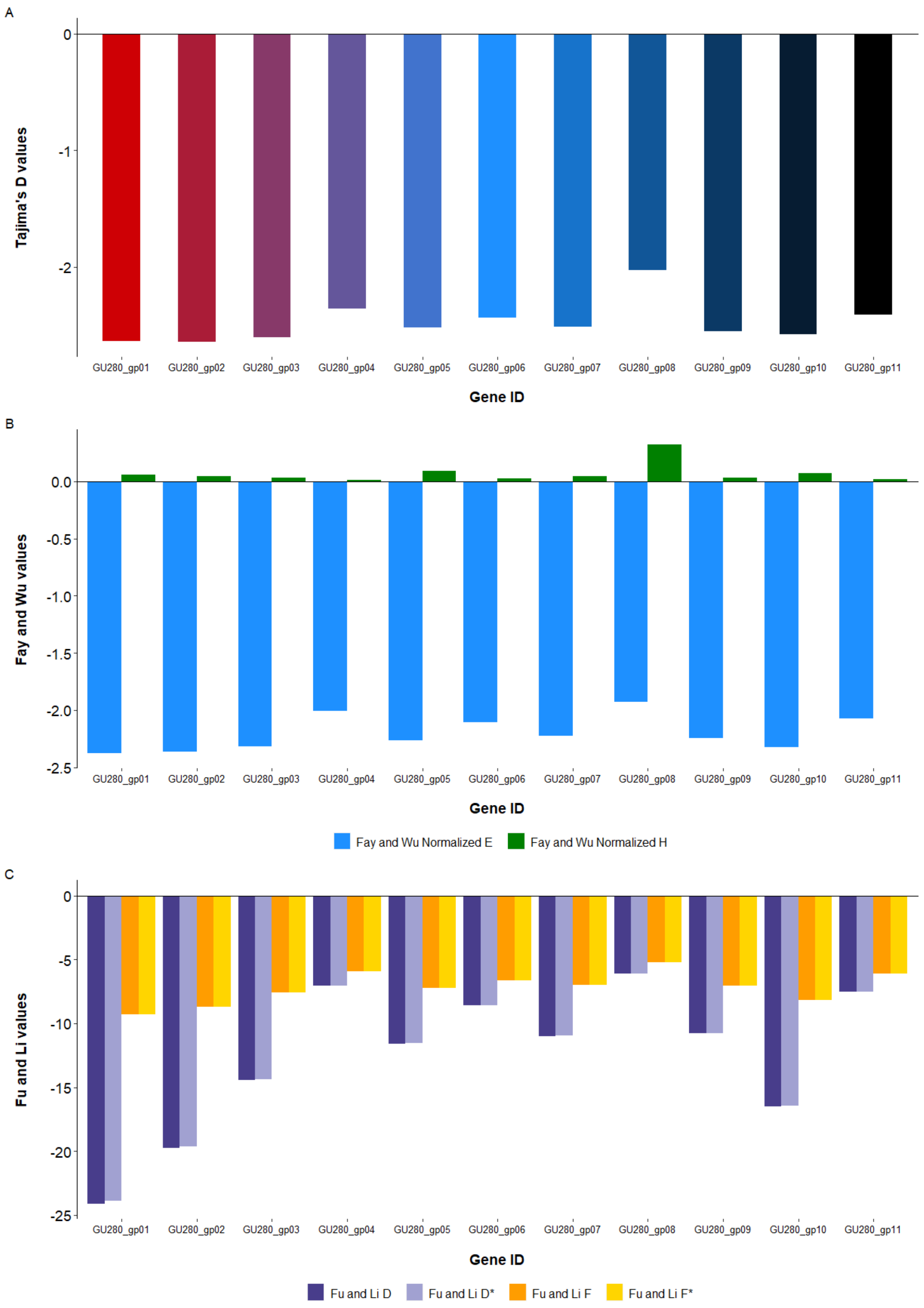

**Figure S - 7.** Graphical representation of the results of the Neutrality test analysis via CATE for 11 genes present in the SARS-CoV-2 viral genome from the Delta strain. (**A**) Depicts the results of the Tajima's D test, (**B**) depicts the results for the Fay and Wu test statistics and (**C**) depicts the results for the Fu and Li test statistics.

#### 5.3. Fixation Index ($F_{st}$)

We have implemented a whole population wide fixation index algorithm in CATE. Most implementations use a pairwise population $F_{st}$ algorithm. This allows us to measure the measures of population variation among multiple populations. CATE's implementation allows the use of unlimited subpopulations.

Therefore, to validate our implementation, it required us to use a test data set on CATE.

##### 5.3.1. Validation of the $F_{st}$ algorithm

Our toy test is based on the following online worked example:
http://www.uwyo.edu/dbmcd/popecol/maylects/fst.html

Our query population structure is as shown in **Table S - 18**. The VCF file structure was custom designed to denote the population structure depicted below.

**Table S - 18.** Detailed overview of the population structure as well as the genotype distributions in each subpopulation.

| | Genotypes | | | Total population (N) |
|---|---|---|---|---|
| | **AA (REF\|REF)** | **Aa (REF\|ALT)** | **aa (ALT\|ALT)** | |
| **Sub population 1** | 125 | 250 | 125 | 500 |
| **Sub population 2** | 50 | 30 | 20 | 100 |
| **Sub population 3** | 100 | 500 | 400 | 1000 |

**1) Allele frequency calculation**

$$\textit{Frequency of allele A in sub population 1} = p_1 = \frac{2 \times 125 + 250}{(500 \times 2)} = 0.50$$

$$\textit{Frequency of allele a in sub population 1} = q_1 = 1 - 0.50 = 0.50$$

$$\textit{Frequency of allele A in sub population 2} = p_2 = \frac{2 \times 50 + 30}{(100 \times 2)} = 0.65$$

$$\textit{Frequency of allele a in sub population 2} = q_2 = 1 - 0.65 = 0.35$$

$$\textit{Frequency of allele A in sub population 3} = p_3 = \frac{2 \times 100 + 500}{(1000 \times 2)} = 0.35$$

$$\textit{Frequency of allele a in sub population 3} = q_3 = 1 - 0.35 = 0.65$$

2) **Estimate the expected heterozygosity for each sub population**

$$H_{exp} = 1 - (p^2 + q^2)$$

| Sub population 1 | Sub population 2 | Sub population 3 |
|---|---|---|
| $H_{exp}1 = 0.5$ | $H_{exp}2 = 0.455$ | $H_{exp}3 = 0.455$ |

3) **Calculate the frequency of allele A (reference) and allele a (alternate) over the total overall population**

$$Frequency\ of\ A = \bar{p} = \frac{(0.5 \times 1000) + (0.65 \times 200) + (0.35 \times 2000)}{3200} = 0.4156$$

$$Frequency\ of\ a = \bar{q} = 1 - 0.4156 = 0.5844$$

4) **Global heterozygosity indices**

**Expected heterozygosity in the sub populations**

$$\boldsymbol{H_S} = \frac{\sum_{1}^{number\ of\ sup\ populations}(H_{exp} \times N_{subpopulation})}{N_{Total}} = 0.4691$$

**Expected heterozygosity in the overall population**

$$\boldsymbol{H_T} = 1 - (\bar{p}^2 + \bar{q}^2) = 0.4858$$

5) **$F_{st}$ value**

$$F_{st} = \frac{H_T - H_S}{H_T} = \frac{0.4858 - 0.4691}{0.4858} = \mathbf{0.0344}$$

The above result was evaluated with the output of CATE given in **Table S - 19**.

**Table S - 19.** CATE's $F_{st}$ output result. The result matches that of the test data proving that the implemented algorithm is correct and works as intended.

| Gene_name | Coordinates | Total_Fst | Total_seg_sites | Average_Fst |
|---|---|---|---|---|
| TEST0 | 01:10504:10506 | 0.034377 | 1 | **0.034377** |

Comparison of the values prove that CATE's $F_{st}$ algorithm works as intended.

#### 5.3.2. $F_{st}$ runtimes

$F_{st}$ function's configurations of all genes and window test types were benchmarked. The individual chromosome runtimes from the 1000 Genomes dataset are shown in **Table S - 20** and those from the 1001 Genomes dataset are depicted in **Table S - 21**.

**Table S - 20.** Total run time taken by CATE to process each chromosome in the 1000 Genomes dataset with a combination of three sub populations of African (AFR), East Asians (EAS), and Europeans (EUR). The results show the run time taken for each test type, with the window test type consisting of both the window size and step size being 10,000 bases.

| Chromosome | Window run time (s) | All genes run time (s) |
|---|---|---|
| 1 | 999 | 523 |
| 2 | 1073 | 471 |
| 3 | 941 | 451 |
| 4 | 795 | 409 |
| 5 | 755 | 345 |
| 6 | 704 | 309 |
| 7 | 678 | 372 |
| 8 | 659 | 382 |
| 9 | 571 | 237 |
| 10 | 554 | 259 |
| 11 | 581 | 291 |
| 12 | 543 | 284 |
| 13 | 388 | 153 |
| 14 | 359 | 207 |
| 15 | 370 | 208 |
| 16 | 373 | 227 |
| 17 | 339 | 238 |
| 18 | 309 | 172 |
| 19 | 294 | 178 |
| 20 | 278 | 144 |
| 21 | 156 | 80 |
| 22 | 154 | 97 |
| **Total runtime (s)** | **11873** | **6037** |
| **Average runtime (s)** | **539.68** | **274.41** |

**Table S - 21.** Total run time taken by CATE to process each chromosome in the 1000 Genomes dataset with a combination of three sub populations of Asia, Western Europe, and Central Europe. The results show the run time taken for each test type, with the window test type consisting of both the window size and step size being 10,000 bases.

| Chromosome | Window run time (s) | All genes run time (s) |
|---|---|---|
| 1 | 28 | 89 |
| 2 | 18 | 38 |
| 3 | 24 | 42 |
| 4 | 18 | 35 |
| 5 | 24 | 48 |
| **Total runtime (s)** | **112** | **252** |
| **Average runtime (s)** | **22.40** | **50.40** |

#### 5.4. Extended Haplotype Homozygosity (EHH)

CATE's implementation of EHH algorithm was validated via the selscan software. All available modes on EHH were tested using the 1000 and 1001 Genomes projects. For FIXED mode the all genes test type was used whereas for the testing of the modes SNP and BP, for each chromosome 500 SNPs were randomly selected. For the benchmarking of CATE's speed, for the 1000 Genomes project dataset we used only the SNPs with MAF greater than 0.05 or common SNPs.

##### 5.4.1. Validation of EHH algorithm

The results of CATE's implementation of EHH were validated using the selscan (8) software and the 1000 Genomes dataset. The SNP rs531730856 located at position 13273 of Chromosome 1 in the Human Genome GRCh37 was used. In CATE's fixed mode it was able to identify the two core haplotypes and separately calculate the EHH values for each haplotype for the given extended region (**Table S - 22**), in the BP and SNP modes CATE was able to detect the extended haplotype regions and calculate their EHH values for both the reference (0) and alternate (1) allele core haplotypes respectively (**Table S - 23**). The values obtained coincided with those from selscan.

**Table S - 22:** Comparison of CATE's results for the EHH test statistic against selscan for the FIXED mode.

| Core haplotype coordinates | Extended haplotype coordinates | Core haplotype number | CATE | selscan |
|---|---|---|---|---|
| 01:13273:13273 | 01:13273:13284 | 1 | 0.996915 | 0.996915 |
| 01:13273:13273 | 01:13273:13284 | 2 | 1 | 1.000000 |

**Table S - 23.** A snippet of the results of CATE's results for the EHH test statistic against selscan from the SNP and BP modes.

| Displacement | SNP position | CATE 0 | selscan 0 | CATE 1 | selscan 1 |
|---|---|---|---|---|---|
| -157 | 13116 | 0.82065 | 0.82065 | 0.852446 | 0.852446 |
| -155 | 13118 | 0.82065 | 0.82065 | 0.852446 | 0.852446 |
| -117 | 13156 | 0.999118 | 0.999118 | 0.995798 | 0.995798 |
| -14 | 13259 | 0.999118 | 0.999118 | 1 | 1.000000 |
| 0 | 13273 | 1 | 1.000000 | 1 | 1.000000 |
| 11 | 13284 | 0.996915 | 0.996915 | 1 | 1.000000 |
| 16 | 13289 | 0.995594 | 0.995594 | 1 | 1.000000 |
| 40 | 13313 | 0.995153 | 0.995153 | 1 | 1.000000 |
| 92 | 13365 | 0.994713 | 0.994713 | 1 | 1.000000 |

#### 5.4.2. All genes test type for FIXED mode

The time taken might vary based on the complexity of the specified core regions. Meaning if the regions have a higher number of unique haplotypes, the processing times can get longer. The results of our test group for the 1000 Genomes dataset are shown in **Table S - 24** and for the 1001 Genomes dataset are shown in **Table S - 25**.

**Table S - 24.** Total run time taken by each chromosome for all genes test type for the 1000 Genomes dataset, including the number of unique core haplotypes that were discovered in each chromosome for the query gene regions that were provided.

| Chromosome | No. unique core haplotypes | Total run time (s) |
|---|---|---|
| 1 | 28266 | 212 |
| 2 | 21858 | 187 |
| 3 | 15380 | 136 |
| 4 | 14876 | 130 |
| 5 | 14336 | 136 |
| 6 | 19479 | 131 |
| 7 | 15767 | 128 |
| 8 | 12378 | 114 |
| 9 | 12310 | 100 |
| 10 | 10283 | 105 |
| 11 | 22101 | 131 |
| 12 | 12074 | 115 |
| 13 | 6827 | 66 |
| 14 | 20181 | 94 |
| 15 | 9273 | 82 |
| 16 | 10021 | 96 |
| 17 | 13962 | 103 |
| 18 | 5394 | 60 |
| 19 | 12261 | 98 |
| 20 | 5750 | 56 |
| 21 | 7011 | 40 |
| 22 | 8855 | 52 |
| **Total** | **298643** | **2372** |
| **Average** | **13574.68** | **107.82** |

**Table S - 25.** Total run time taken by each chromosome for all genes test type for the 1001 Genomes dataset, including the number of unique core haplotypes that were discovered in each chromosome for the query gene regions that were provided.

| Chromosome | No. unique core haplotypes | Total run time (s) |
|---|---|---|
| 1 | 31986 | 56 |
| 2 | 38714 | 34 |
| 3 | 81897 | 41 |
| 4 | 37986 | 48 |
| 5 | 31571 | 73 |
| **Total** | **222154** | **252** |
| **Average** | **44430.8** | **50.40** |

#### 5.4.3. SNP and BP modes

For each chromosome 500 SNPs were selected at random. In SNP mode a window of 250 SNPs was used and in BP mode a window of 100000 base pairs was used to define the extended haplotype region on each side the core haplotype SNP. The results of the 1000 Genomes dataset are depicted in **Table S - 26** and for the 1001 Genomes dataset is depicted in **Table S - 27**.

**Table S - 26.** Total run time taken by each chromosome for the 1000 Genomes dataset to process 500 SNPs per chromosome in both BP and SNP modes in the EHH function.

| Chromosome | BP mode run time (s) | SNP mode run time (s) |
|---|---|---|
| 1 | 3143 | 2110 |
| 2 | 2757 | 1766 |
| 3 | 2906 | 1630 |
| 4 | 3023 | 1579 |
| 5 | 2416 | 1674 |
| 6 | 5109 | 1544 |
| 7 | 3271 | 1856 |
| 8 | 3054 | 1639 |
| 9 | 3744 | 2131 |
| 10 | 3576 | 2016 |
| 11 | 3165 | 1759 |
| 12 | 3245 | 1928 |
| 13 | 3169 | 1760 |
| 14 | 4375 | 2043 |
| 15 | 3784 | 2333 |
| 16 | 5415 | 2520 |
| 17 | 4506 | 2642 |
| 18 | 3738 | 2120 |
| 19 | 5300 | 2646 |
| 20 | 3907 | 2370 |
| 21 | 4166 | 2313 |
| 22 | 6177 | 3015 |
| **Total runtime (s)** | **83946** | **45394** |
| **Average runtime (s)** | **3815.73** | **2063.36** |

**Table S - 27.** Total run time taken by each chromosome for the 1001 Genomes dataset to process 500 SNPs per chromosome in both BP and SNP modes in the EHH function.

| Chromosome | BP mode run time (s) | SNP mode run time (s) |
|---|---|---|
| 1 | 274 | 531 |
| 2 | 479 | 344 |
| 3 | 247 | 213 |
| 4 | 408 | 334 |
| 5 | 246 | 520 |
| **Total runtime (s)** | **1654** | **1942** |
| **Average runtime (s)** | **330.80** | **388.40** |

#### 5.5. McDonald–Kreitman (MK) Neutrality Index (NI)

Due to the complexities of the MK test and the unique assumptions that have had to be made due to the use of VCF formats, our implementation of CATE was validated using a toy example.

##### 5.5.1. Validation of the MK test

We first took a known Open Reading Frame (ORF) region. The candidate region was from the Cathepsin E (CTSE) protein on chromosome 1. The region lies within the base pairs of 206,330,863 and 206,331,228 of chromosome 1 of the human genome reference GRCh37 (33, 34).

The reference sequence is as follows:

ATGAAGCAGGAGGAAGAACCCACCGACCAAAATGGTGGAACCCACCAGCTGTCCAAGGCCACC
CAGTGCCAGCCACCACCACCACACAGGCAGCCTCCAGGCAGTACTACTACCTTCTTCCCCCGTT
TGTCATTTGCATCCTTCTTTCCAACCCACAGGACTTCGTGGATGGAATGCAGTTCTGCAGCAGTG
GCTTTCAAGGACTTGACATCCACCCTCCAGCTGGGCCCCTCTGGATCCTGGGGGATGTCTTCAT
TCGACAGTTTTACTCAGTCTTTGACCGTGGGAATAACCGTGTGGGACTGGCCCCAGCAGTCCCC
TAAGGAGGGGCCTTGTGTCTGTGCCTGCCTGTCTGACAGACCTTGA

The translated amino acid sequence:

MKQEEEPTDQNGGTHQLSKATQCQPPPPHRQPPGSTTTFFPRLSFASFFPTHRTSWMECSSAAVA
FKDLTSTLQLGPSGSWGMSSFDSFTQSLTVGITVWDWPQQSPKEGPCVCACLSDRPX

The outgroup sequence was selected from the mouse (*Mus musculus*). They too hold the CTSE gene in their chromosome 1. The region of the outgroup sequence that aligned with the reference was:

AGGACTTGACATTCCACCTCCAGCTGGGCCCCTCTGGATCCTGGGGGATGTCTTCATCCGACAG
TTCTACTCAGTCTTTGACCGTGGAAATAACCAAGTGGGATTGGCCCCCGCAGTTCCCTAAAGAG
GG

The alignment of the reference with the outgroup was as follows, **Figure S - 8**:

```
Query  200  AGGACTTGACA-TCCACCCTCCAGCTGGGCCCCTCTGGATCCTGGGGGATGTCTTCATTC  258
            ||||||||||| |||| ||||||||||||||||||||||||||||||||||||||||| |
Sbjct  1    AGGACTTGACATTCCA-CCTCCAGCTGGGCCCCTCTGGATCCTGGGGGATGTCTTCATCC  59

Query  259  GACAGTTTTACTCAGTCTTTGACCGTGGGAATAACCGTGTGGGACTGGCCCCAGCAGTCC  318
            ||||||| |||||||||||||||||||| |||||||  |||||| ||||||| ||||| |
Sbjct  60   GACAGTTCTACTCAGTCTTTGACCGTGGAAATAACCAAGTGGGATTGGCCCCCGCAGTTC  119

Query  319  CCTAAGGAGGG  329
            ||||| |||||
Sbjct  120  CCTAAAGAGGG  130
```

**Figure S - 8.** Capture of the NCBI’s blastn alignment format of the human reference (Query) sequence with that of the mouse (Sbjct) sequence.

A subset of the population was then altered to hold the 13 mutations whose details are shown in **Table S - 28**.

**Table S - 28.** Positions of all 13 simulated mutations and the alternate alleles they occupy.

| Number | Position | Alternate allele |
|---|---|---|
| 1 | 206330886 | T |
| 2 | 206330953 | G |
| 3 | 206330964 | A |
| 4 | 206330976 | T |
| 5 | 206330986 | T |
| 6 | 206331008 | C |
| 7 | 206331027 | A |
| 8 | 206331092 | A |
| 9 | 206331101 | T |
| 10 | 206331120 | T |
| 11 | 206331169 | T |
| 12 | 206331190 | A |
| 13 | 206331193 | T |

Not all 13 positions will align with the outgroup sequence. When aligned with the outgroup sequence 11 codons were identified. The details of the positions of these codons are described in **Table S - 29**.

**Table S - 29.** Description of the 11 codons and the positions of their triads in the genome.

| Codon number | Position 1 | Position 2 | Position 3 |
|---|---|---|---|
| 1 | 206331091 | 206331092 | 206331093 |
| 2 | 206331100 | 206331101 | 206331102 |
| 3 | 206331118 | 206331119 | 206331120 |
| 4 | 206331127 | 206331128 | 206331129 |
| 5 | 206331148 | 206331149 | 206331150 |
| 6 | 206331157 | 206331158 | 206331159 |
| 7 | 206331163 | 206331164 | 206331165 |
| 8 | 206331169 | 206331170 | 206331171 |
| 9 | 206331172 | 206331173 | 206331174 |
| 10 | 206331178 | 206331179 | 206331180 |
| 11 | 206331184 | 206331185 | 206331186 |

**Table S - 30.** Detailed overview of the 11 codons that align with the outgroup regions and the population samples. Bases that have been subject to substitutions in the reference sequence are indicated in green. The SNPs that are fixed substitutions are demarcated in black for nonsynonymous mutations ($D_n$) and maroon for synonymous substitutions ($D_s$). The within species polymorphisms are denoted by grey and orange colours for nonsynonymous ($P_n$) and synonymous substitutions ($P_s$) respectively. As shown the substitution count is calculated by a parsimonious method, meaning they are all direct single base mutations.

| Codon number | 01 | 02 | 03 | 04 | 05 | 06 | 07 | 08 | 09 | 10 | 11 |
|---|---|---|---|---|---|---|---|---|---|---|---|
| Reference (Ref.) | C C C | T C C | T T C | T T T | G G A | G T G | G A C | C C C | C A G | T C C | A A G |
| Ref. amino acid | P | S | F | F | G | V | D | P | Q | S | K |
| Outgroup (Out.) | C C C | T C C | T C C | T C T | G A A | A A G | G A T | C C C | C C G | T T C | A A A |
| Out. amino acid | P | S | S | S | E | M E | D | P | P | F | K |
| Sample | C A C | T C T | T T T | T T T | G G A | G T G | G A C | T C C | C A G | T C C | A A G |
| Sample amino acid | H | F | F | F | G | V | D | S | Q | S | K |
| Dn Total: 7 | | | 1 | 1 | 1 | 2 | | | 1 | 1 | |
| Ds Total: 2 | | | | | | | 1 | | | | 1 |
| Pn Total: 3 | 1 | 1 | | | | | | 1 | | | |
| Ps Total: 1 | | | 1 | | | | | | | | |

$$Neutrality\ Index\ (NI) = \frac{P_n/P_s}{D_n/D_s} = \frac{3/1}{7/2} = \frac{3}{3.5} = \mathbf{0.8571429}$$

The above result was evaluated with the output given by CATE in **Table S - 31**.

**Table S - 31.** CATE's output for the MK test. Comparison of the values with that of our example show that the algorithm works as intended.

| Gene_name | Gene_Coordinates | ORF_Coordinates | Ds | Dn | Ps | Pn | Neutrality_Index |
|---|---|---|---|---|---|---|---|
| TEST0 | 01:<br>206330863:<br>206331228 | 01:<br>206330863:<br>206331228 | **2** | **7** | **1** | **3** | **0.857143** |

The comparison of the $D_n$, $D_s$, $P_n$ and $P_s$ values and the NI result with that of CATE's results show that the algorithm is working correctly. CATE was configured with ORF known mode set to "YES".
